# MiR-208b-3p is an upstream regulator of redox-associated cardiomyocyte survival

**DOI:** 10.64898/2026.10.20.760401

**Authors:** Mohammad Bakhtiar Hossain, Humam Siga, Zaher ElBeck

## Abstract

**Background:** Cardiac stress induces *MYH6*-to-*MYH7* remodeling and activates redox-defense programs, but the regulatory mechanisms that distinguish adaptive from maladaptive stress responses remain incompletely defined. We investigated the role of miR-208b-3p, an intronic *MYH7* microRNA, in cardiac stress adaptation.

**Methods and Results:** We integrated myocardial transcriptomic and small RNA sequencing from stress-adapted *Mlp^−/−^* mice and human cardiomyopathy to identify stress-adaptive programs associated with preserved myocardial viability and complemented these analyses with *in vivo* miR-208b-3p inhibition and cellular gain-of-function experiments. *Mlp^−/−^*hearts exhibited marked *Myh7* induction accompanied by an ≈20-fold increase in miR-208b-3p, whereas this coordinated response was blunted in human end-stage dilated and ischemic cardiomyopathies. *In vivo* inhibition of miR-208b-3p selectively reduced *Myh7*, without altering *Myh6* or baseline cardiac function. This reduction in *Myh7* occurred without induction of established *Myh7* transcriptional repressors, and miR-208b-3p gain-of-function similarly reduced *MYH7* while suppressing several canonical repressors, indicating that its relationship with *MYH7* extends beyond the established repressor pathway. miR-208b-3p inhibition activated oxidative stress, NRF2, autophagy, and cell-survival pathways. Oxidative stress transiently suppressed miR-208b-3p, with rebound induction after stress removal, whereas antioxidant treatment increased its expression. Pharmacological NRF2 activation did not alter miR-208b-3p, whereas miR-208b-3p gain-of-function reduced oxidative stress-associated apoptotic signaling and suppressed ATM, a predicted target and regulator of cellular stress responses, supporting an upstream role for miR-208b-3p in stress adaptation.

**Conclusions:** MiR-208b-3p is a dynamic component of cardiac stress adaptation that links myosin remodeling with redox and survival responses. Its regulation of ATM and association with NRF2-related stress responses support an upstream role in coordinating cellular adaptation to cardiac stress.

**Graphical abstract:** 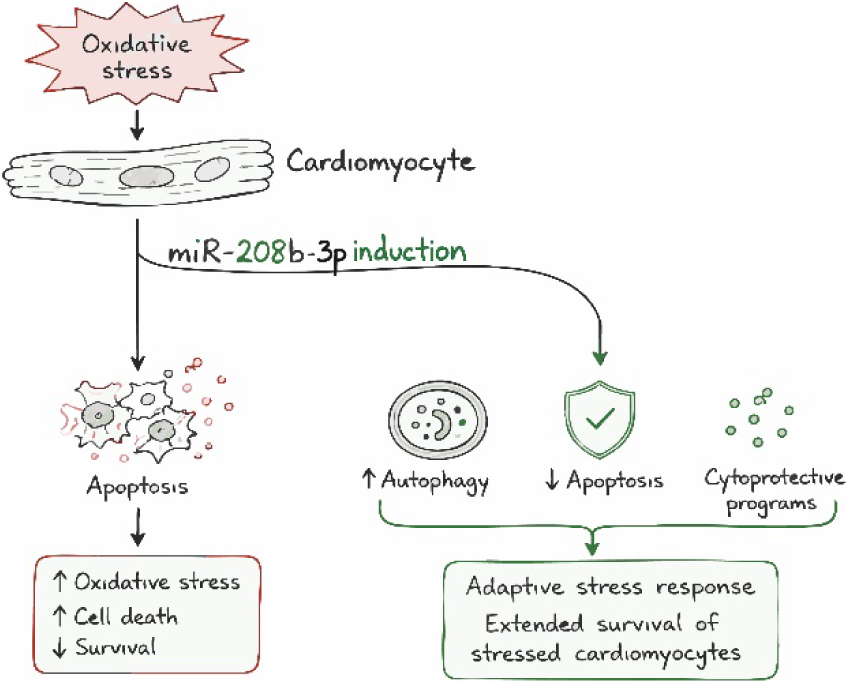

## Introduction

MicroRNAs (miRNAs) are central post-transcriptional regulators of cardiac development, stress adaptation, and pathological remodeling in heart failure. Among these, the muscle-specific myomiR family, comprising miR-208a-3p, miR-208b-3p, and miR-499-5p, has classically been defined by its role in regulating the reciprocal expression of the cardiac myosin heavy chain (MHC) genes, *MYH6* (*α*-MHC) and *MYH7* (*β*-MHC) (van Rooij, Quiat et al. 2009). Although *α*-MHC and *β*-MHC share 93% sequence homology, *α*-MHC possesses significantly higher actin-attachment and cross-bridge kinetics. In the failing human heart, contraction-relaxation dysfunction is accompanied by a shift from *α*-MHC to *β*-MHC, characteristic of a fetal gene reprogramming profile (Palmer 2005).

Intriguingly, despite sharing identical 5′-seed sequences, members of the myomiR family exhibit distinct clinical and functional profiles. The expression of miR-208a-3p correlates with myocardial fibrosis and adverse clinical outcomes in dilated cardiomyopathy (DCM) (Satoh, Minami et al. 2010). Conversely, circulating levels of miR-208b-3p spike dramatically (over 1,000-fold) following acute myocardial infarction (Corsten, Dennert et al. 2010, Gidlof, Andersson et al. 2011), coinciding with a surge in myocardial oxidative stress (Grieve, Byrne et al. 2004, Hori and Nishida 2009). Furthermore, dynamic shifts in myomiR expression have been observed in DCM patients undergoing reverse remodeling in response to beta-blocker therapy (Sucharov, Kao et al. 2017), suggesting that these microRNAs actively track changes in cardiac stress and functional recovery.

Cardiomyocytes maintain redox homeostasis through an integrated network of antioxidants, transcriptional regulators, and redox-sensitive miRNAs termed "redoximiRs" (Cheng, Ku et al. 2013, Fierro-Fernandez, Miguel et al. 2016). RedoximiRs govern intracellular redox balance by modulating key node proteins, including the master antioxidant transcription factor NRF2 (Cheng, Ku et al. 2013, Kurinna and Werner 2015), as well as reactive oxygen species (ROS)-generating enzymes (Manea, Constantin et al. 2015). Reciprocally, cellular redox status regulates miRNA biogenesis and transcriptional activity through oxidative modifications and epigenetic remodeling (Carbonell and Gomes 2020).

We previously demonstrated that the left ventricular myocardium of patients with end-stage DCM shares gene expression signatures related to mitochondrial dysfunction and altered oxidative phosphorylation with muscle LIM protein deficient (*Mlp*^-/-^ or *Csrp3*^-/-^) mice, a model of eccentric hypertrophy and DCM (Arber, Hunter et al. 1997, ElBeck, Hossain et al. 2024). However, these systems differ markedly in phenotypic trajectory. While human end-stage DCM myocardium is characterized by terminal heart failure and cellular exhaustion, *Mlp^-/-^* mice display notable survival resilience, engaging compensatory bioenergetic responses and reverse cardiac remodeling, particularly in females (ElBeck, Hossain et al. 2024). Nonetheless, clinical observations demonstrate that patients with end-stage DCM can undergo substantial reverse remodeling and functional recovery following mechanical unloading via left ventricular assist device (LVAD) support (Ambardekar and Buttrick 2011, Burkhoff, Topkara et al. 2021). This indicates that latent cardioprotective programs remain accessible even in failing human myocardium. Uncovering the regulatory mechanisms that differentiate the adaptive state in mice models from terminal human DCM may illuminate endogenously protective pathways that preserve cardiomyocyte viability under chronic strain.

To uncover these regulatory mechanisms, we performed comparative cross-species transcriptomic screening, which identified a blunted miR-208b-3p response and suppressed downstream survival architectures in end-stage human DCM compared to robust induction in *Mlp*^-/-^ hearts. Mechanistically, we show that miR-208b-3p acts upstream of the NRF2 axis by targeting the ROS sensor *ATM*, functioning as a temporal survival brake that suppresses pro-apoptotic drivers like *Osgin1* under oxidative strain. These findings redefine miR-208b-3p not merely as a passive marker of myosin gene expression, but as an upstream homeostatic determinant of cardiomyocyte survival under bioenergetic and oxidative strain.

## Results

### Distinct and shared myocardial transcriptomic signatures between human DCM and *Mlp⁻/⁻* mice

To identify endogenous protective mechanisms that drive adaptive remodeling in *Mlp⁻/⁻* mice but are lacking or suppressed in end-stage human dilated cardiomyopathy, we intersected transcriptomic profiles from human left ventricular (LV) myocardium (harboring mutations in *LMNA*, *RBM20*, or *TTN*) (Sielemann, Elbeck et al. 2020) with LV tissue from the survival-resilient murine DCM *Mlp⁻/⁻* model (ElBeck, Hossain et al. 2024) (**Figure 1A**). This comparative myocardial transcriptomic analysis revealed a shared signature of 316 differentially expressed genes (DEGs), alongside substantial distinct gene programs: human end-stage DCM hearts exhibited 1,034 uniquely dysregulated genes, where *Mlp⁻/⁻* hearts exhibited 2,800 unique DEGs (**Figure 1A**).

**Figure 1:**
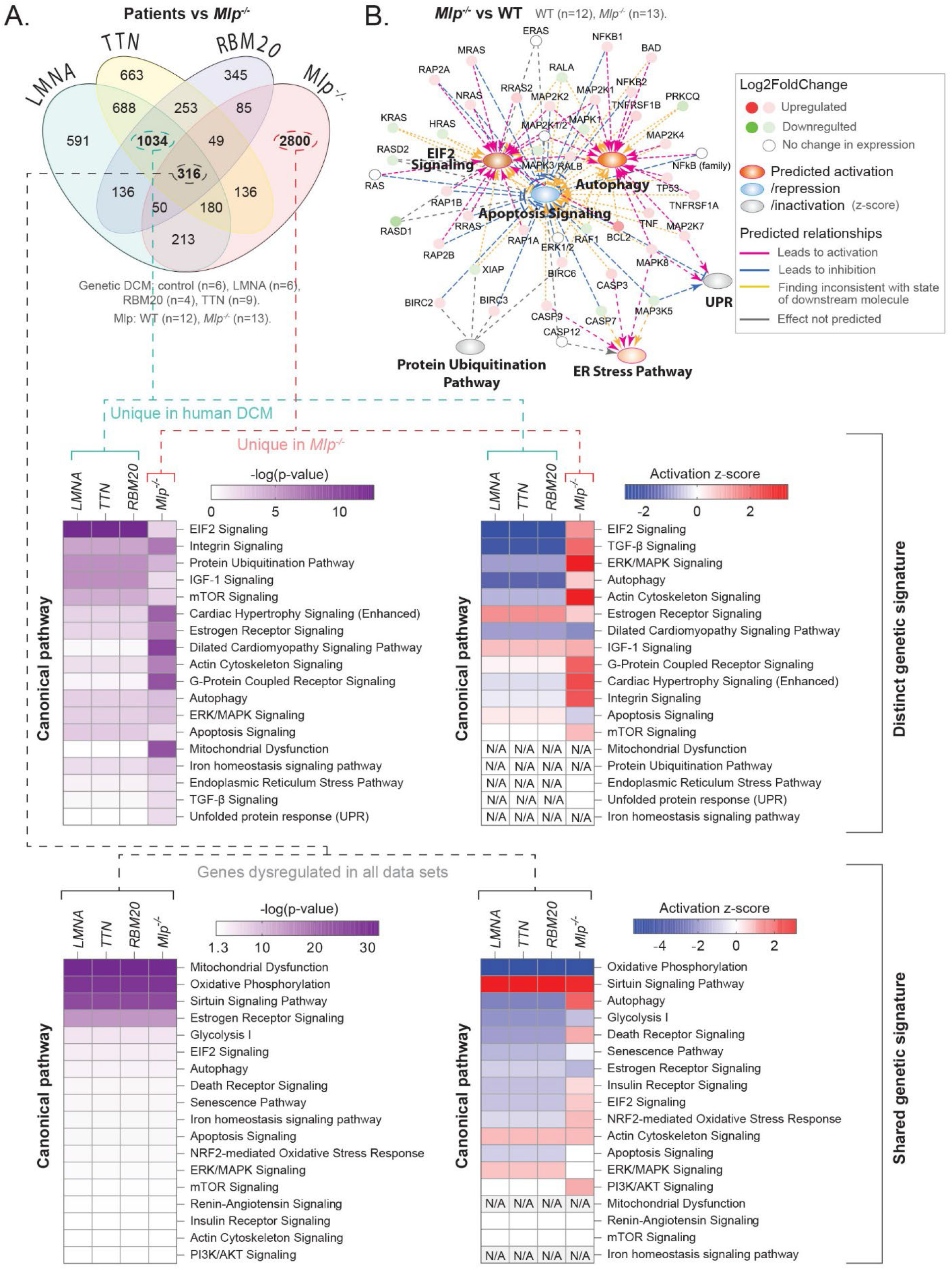
Shared and distinct genetic signatures between human and murine genetic-DCMs*^-^*. **A.** Venn diagram of differentially expressed genes in genetic DCM obtained through intersecting transcriptomic data on LV myocardium of patients with DCM (Sielemann, Elbeck et al. 2020) and male and female *Mlp^-/-^ (ElBeck, Hossain et al. 2024)*.Only DEGs with *p<0.05* and adjusted *P* value < 0.01 were included in this analysis. Lower panels: Selected pathways from IPA enrichment analysis on DEGs that were included in the intersections, highlighted with bold font in the Venn diagram. **B.** IPA *in silico* map that connects some differential expressed genes of interest with phenotype utilizing public studies. Indicated pathways were selected from figure A1 based on their relation to ER stress. The map was then overlayed to the transcriptomic data on myocardium from *Mlp^-/-^* mice described in figure A1.

Ingenuity Pathway Analysis (IPA) revealed that the shared transcriptomic signature between human and murine DCM primarily comprises genes encoding mitochondrial and oxidative phosphorylation machinery (**Figure 1A**, bottom panel). In contrast, genes uniquely dysregulated in either end-stage human DCM or *Mlp⁻/⁻* hearts mapped to multiple pathways with opposing activation trends, including eIF2 signaling, TGF-β signaling, and autophagy (**Figure 1A(**upper panel **) and 1B**), several of which are associated with endoplasmic reticulum (ER) stress and protein misfolding. ER stress can arise from disturbances in calcium homeostasis and oxidative stress, leading to the accumulation of misfolded proteins in the ER lumen and activation of the unfolded protein response (UPR). To restore ER homeostasis, cells rely on protein-degradation systems, including the proteasome and autophagy. Although these mechanisms initially help cells adapt to stress, severe or prolonged stress that exceeds their capacity can promote apoptotic cell death (Sozen, Karademir et al. 2015). Thus, rather than progressing toward cell death, the *Mlp*^⁻/⁻^ myocardium appeared to maintain a balance between pro- and anti-apoptotic regulators within these pathways (**Figure 1B**)

This protective stress response in *Mlp⁻/⁻* mice operated alongside classic structural and metabolic adaptations. Under chronic biomechanical and oxidative strain, murine cardiomyocytes reactivate a fetal gene program characterized by a shift from MYH6 to MYH7 (*α*- to *β*-myosin heavy chain), an isoform transition that lowers cross-bridge ATP consumption rate to preserve energy availability under strain (Hasenfuss, Mulieri et al. 1991). Correspondingly, *Mlp⁻/⁻* hearts exhibited a pronounced induction of *Myh7* alongside *Myh6* downregulation (**Figure 2A**), reflecting an active bioenergetic adaptation in the mouse model; in contrast, these isoform shifts were minimal in human DCM, where the vast majority of adult ventricular myocardium natively consists of *β*-MHC at baseline (**Figure 2B, and Supplementary Fig. 2A**).

**Figure 2.**
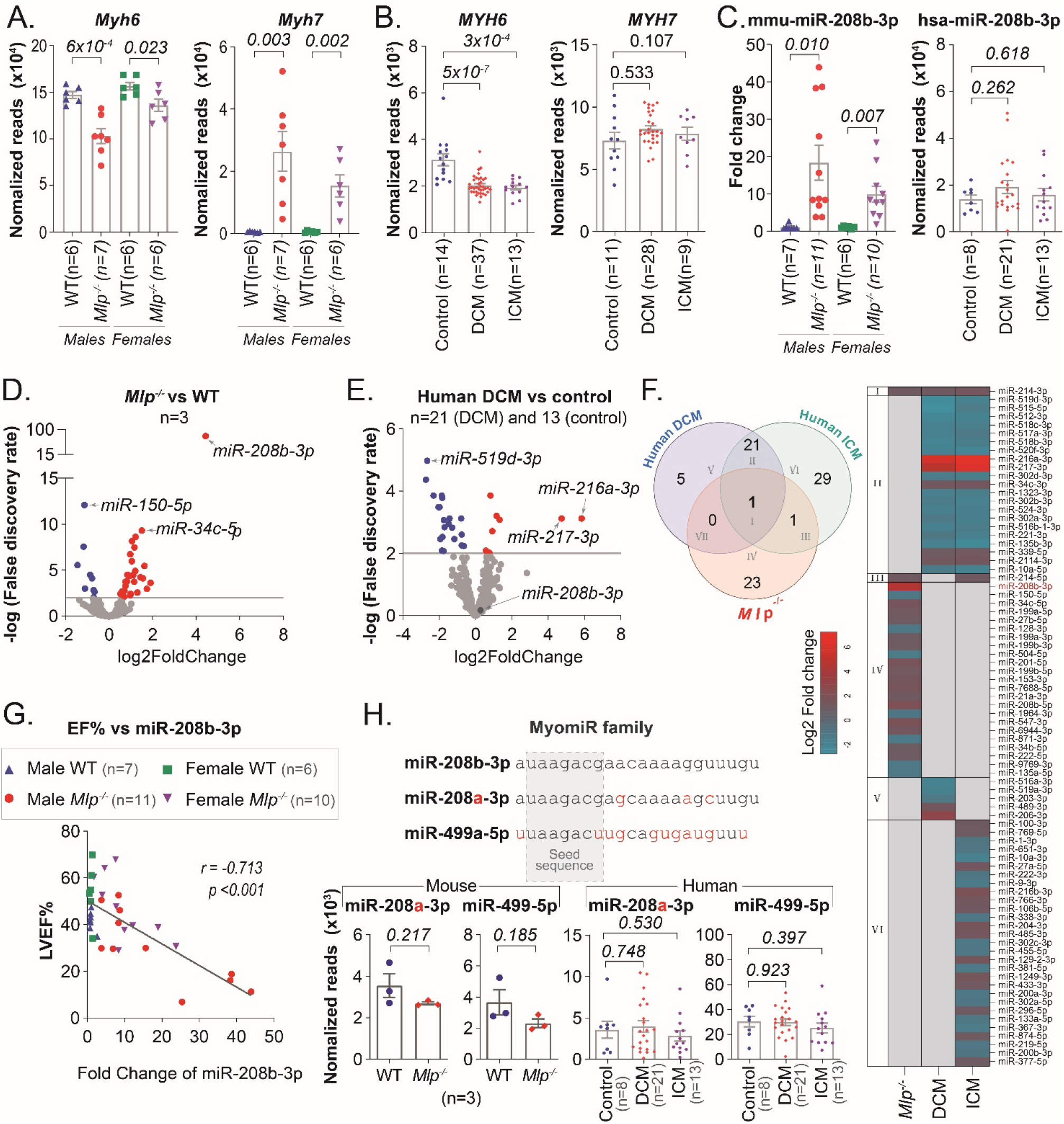
Differential expression of MYH isoforms and miR-208b-3p in *Mlp^−/−^* myocardium and human cardiomyopathy. **A.** *Myh6* and *Myh7* expression in male and female WT and *Mlp^−/−^* mouse myocardium by RNA-seq. **B.** *MYH6* and *MYH7* expression in human control, DCM, and ICM myocardium by RNA-seq. Sample sizes for MYH7 reflect exclusion of invalid data points present in the public GEO dataset (see Methods for details) **C.** miR-208b-3p expression in WT and *Mlp^−/−^* mouse myocardium by qPCR and in human DCM and ICM myocardium by small RNA-seq. **D,E.** Volcano plots of differentially expressed miRNAs in *Mlp^−/−^*versus WT mouse myocardium (D) and DCM versus control human myocardium (E). **F.** Venn diagram and heatmap showing overlapping differentially expressed miRNAs in *Mlp^−/−^*, DCM, and ICM. Differentially expressed miRNAs were defined as FDR < 0.05 and |log2FC| ≥ 1. Heatmap values represent log2FC. **G.** Pearson correlation between LVEF and miR-208b-3p expression in WT and *Mlp^−/−^* myocardium. **H.** Expression of miR-208a-3p and miR-499-5p in WT and *Mlp^−/−^* myocardium and in human control, DCM, and ICM myocardium by small RNA-seq. Data in A and B were adapted from ElBeck et al. (ElBeck, Hossain et al. 2024) and Sweet et al. (Sweet, Cocciolo et al. 2018), respectively. Human small RNA-seq data were adapted from Akat et al. (Akat, Moore-McGriff et al. 2014). Data are mean ± SEM; individual points represent individual samples. Statistical analysis was performed using unpaired two-tailed *t-*tests or Pearson correlation as indicated.

### Selective and pronounced upregulation of miR-208b-3p in *Mlp^-/-^* myocardium

To understand the regulatory network accompanying the marked *Myh7* upregulation in *Mlp^-/-^* hearts, we examined the MyomiR family, specifically miR-208b-3p, an intronic miRNA co-transcribed with *Myh7* that fine-tunes cardiac myosin isoform expression by directly targeting its repressors (van Rooij, Quiat et al. 2009). Profiling microRNA expression across *Mlp^-/-^* hearts as well as human dilated cardiomyopathy (DCM) and ischemic cardiomyopathy (ICM) myocardium revealed that, paralleling the marked *Myh7* induction, mature miR-208b-3p was strongly elevated approximately 20-fold in *Mlp^-/-^* hearts compared with wild-type littermate controls (**Figures 2C and D**). In contrast, this pronounced induction was absent in human DCM/ICM myocardium (**Figures 2C and E**), where the increase in *MYH7* was also comparatively modest (**Figure 2B**).

Comparison between disease phenotypes revealed a substantial miRNA response shared between human DCM and ICM, with 21 miRNAs dysregulated in both human etiologies but absent in *Mlp^-/-^*hearts. Notably, this shared human signature is broadly involved in cardiovascular stress, vascular repair, fibrosis, and cardiac remodeling. In contrast, only miR-214-3p, a known master regulator of cardiac stress adaptive response and calcium handling (Aurora, Mahmoud et al. 2012), was common across all three groups, while several miRNAs uniquely altered in *Mlp^-/-^* hearts have been implicated in cardiac stress resilience (**Figure 2F**). The lack of differential miR-208b-3p expression in terminal human DCM or ICM—despite its marked elevation in survival-adapted *Mlp^-/-^* hearts—highlights miR-208b-3p as a central component of the unique adaptive response that preserves viable myocardium under chronic strain.

Myocardial miR-208b-3p levels strongly and negatively correlated (*r* = −0.713, *p* < 0.001) with left ventricular ejection fraction (LVEF%) across both male and female *Mlp^-/-^* mice (**Figure 2G**), tracking directly with phenotype severity. Conversely, miR-208b-5p—originating from the 5p arm of the same pre-miR-208b—exhibited very low overall expression. While miR-208b-5p was upregulated in *Mlp^-/-^*hearts (**Supplementary figure S2B**), its abundance was negligible compared to miR-208b-3p (∼10 reads vs. ∼20,000 reads). This vast difference indicates that miR-208b-5p acts primarily as a passenger strand subject to expected degradation upon miR-208b-3p loading onto the AGO complex, an asymmetry similarly observed for human miR-208b-3p/5p (**Supplementary fig. S2B**).

Other MyomiR family members sharing the same seed sequence, namely miR-208a-3p and miR-499-5p, showed no significant upregulation in *Mlp^-/-^* or human DCM/ICM hearts (**Figure 2H**), demonstrating that this response is specific to miR-208b-3p rather than a global activation of the MyomiR family.

Notably, the most significantly downregulated microRNA in *Mlp^-/-^*hearts was miR-150-5p (Supplementary **Fig S2C**). MiR-150-5p has been reported to target the epigenetic regulators *EZH1* and *EZH2* (Selvam, Bandi et al. 2022), and EZH2 has been suggested to interact with miR-208b to regulate cardiac *Myh7* expression (Mathiyalagan, Okabe et al. 2014). However, despite high baseline expression and a 2-fold reduction of miR-150-5p in *Mlp^-/-^* myocardium, we observed no corresponding increase in *Ezh1* or *Ezh2* transcript levels (Supplementary **Fig S2D**).

### Knockdown of miR-208b-3p reduces *Myh7* expression and modulates myocardial redox signaling without altering cardiac function in *Mlp^-/-^* mice

Although elevated circulating or tissue miR-208b-3p has not been consistently linked to adverse clinical outcomes in human DCM (Satoh, Minami et al. 2010), its strong inverse correlation with ejection fraction in *Mlp^-/-^* mice (**Figure 2G**) led us to test whether elevated miR-208b-3p actively drives cardiac dysfunction. Male *Mlp^-/-^* mice were treated with LNA-modified miR-208b-3p antagomirs under a one-month period (**Figure 3A**). Despite achieving a >70% reduction in myocardial miR-208b-3p levels (**Figure 3B**), antagomir treatment did not alter cardiac function, left ventricular chamber dimensions, or wall thickness (**Figure 3C–D**). However, miR-208b-3p inhibition induced a selective downregulation of *Myh7* expression without affecting *Myh6* transcript levels (**Figure 3E and Supplementary Figure S3A**).

**Figure 3:**
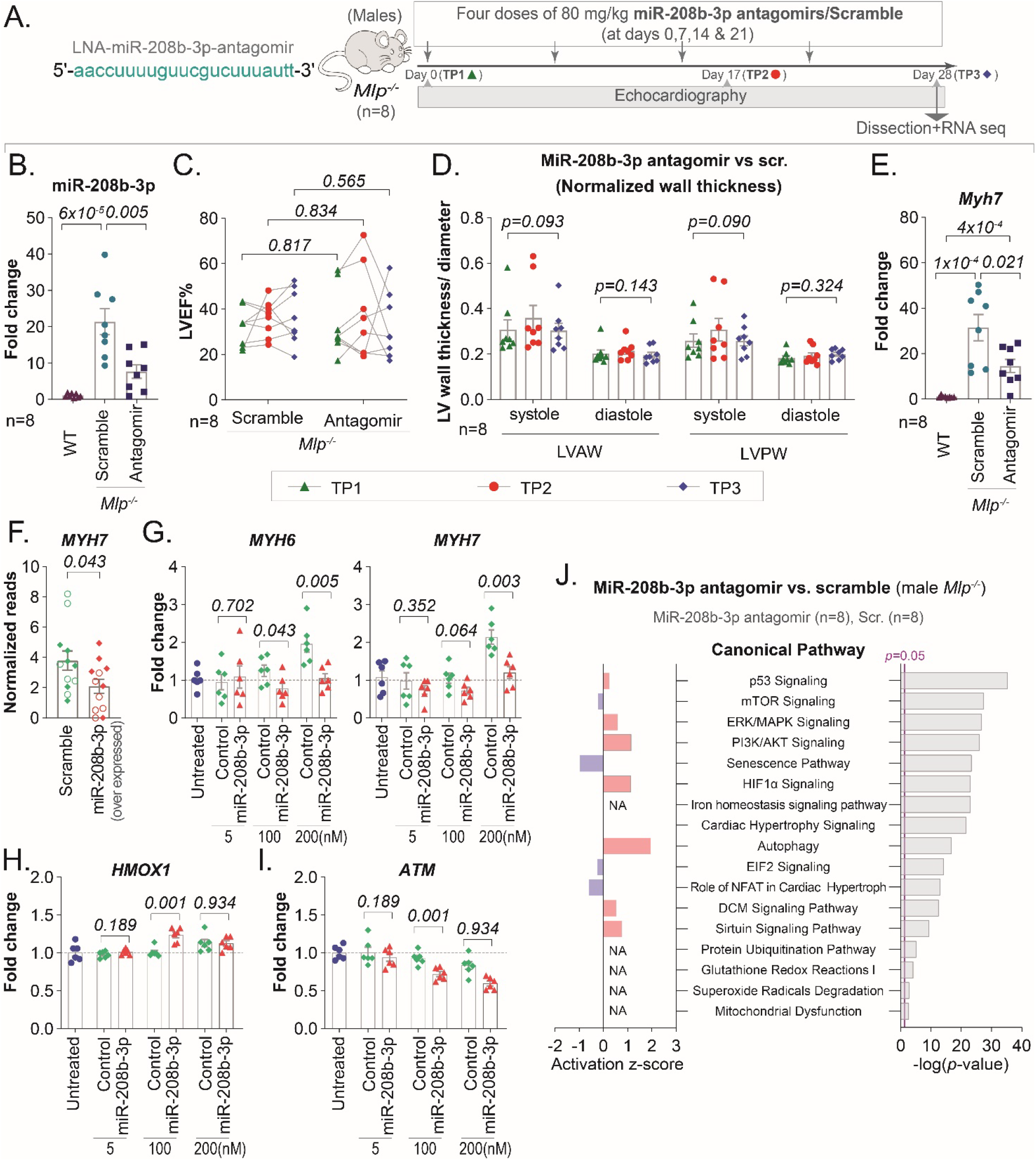
*In vivo* inhibition of miR-208b-3p reduces *Myh7* expression but does not improve cardiac function. **A.** Experimental design for treatment of male *Mlp^-/-^* mice with LNA-modified miR-208b-3p antagomirs or scramble control (80 mg/kg on days 0, 7, 14, and 21). **B.** miR-208b-3p expression in LV myocardium following antagomir treatment, normalized to untreated WT littermates. **C.** Longitudinal changes in LVEF in *Mlp^-/-^* mice treated with miR-208b-3p antagomirs or scramble control. **D.** Longitudinal changes in LVAW and LVPW thickness normalized to LV diameter. **E.** *Myh7* expression in LV myocardium following miR-208b-3p antagomir treatment, normalized to untreated WT littermates. **F.** *MYH7* expression in HEK293 cells overexpressing miR-208b-3p or scramble control, estimated by RNA seq. Filled and open data points represent two independent experimental replicates. **G–I**. *MYH6*, *MYH7*, *HMOX1*, and *ATM* expression in HEK293 cells treated with 5, 100, or 200 nM miR-208b-3p mimic or concentration-matched negative control mimic. **J.** Selected canonical pathways identified by Ingenuity Pathway Analysis of LV transcriptomic data from miR-208b-3p antagomir-treated *Mlp^-/-^* mice. Data are presented as mean ± SEM. Statistical analysis was performed using unpaired two-tailed *t*-tests for **B** and **E–I** and paired two-tailed *t*-tests for **C** and **D**.

To evaluate whether this concurrent decrease in miR-208b-3p and *Myh7* operates through canonical regulation of known *Myh7* transcriptional repressors (*Sox6*, *Purb*, *Cbx1*, *Sp3*, and *Med13*) (van Rooij, Quiat et al. 2009), we examined their expression in treated myocardium. Although antagomir treatment significantly reduced *Myh7* expression in *Mlp^-/-^* hearts (**Figure 3E**), it failed to elevate any of these established repressors (**Supplementary Figure 3B**). Furthermore, none of these repressors were suppressed in baseline *Mlp^-/-^* hearts despite marked upregulation of both miR-208b-3p and *Myh7* (**Figures 2A and 2C**). This discrepancy was mirrored in human DCM myocardium, where most repressors were significantly downregulated despite only modest miR-208b-3p or *MYH7* expression (**Supplementary Figure 3B and Figures 2B-C**).

To evaluate whether miR-208b-3p regulates *MYH7* through direct repression of canonical *MYH7* transcriptional repressors, we performed *in vitro* gain-of-function studies in HEK293 cells using both plasmid vector overexpression and synthetic miRNA mimic transfections. Initial plasmid-driven overexpression of miR-208b-3p significantly decreased *MYH7* transcript levels (**Figures 3C and supplementary figure S3C**); however, this downregulation occurred without altering transcript levels of the canonical repressors (*SOX6*, *PURB*, *CBX1*, *SP3*, and *MED13*; **Supplementary Figure S3D**).

To more precisely isolate the direct post-transcriptional effects of mature miR-208b-3p and eliminate vector-induced cellular stress, cells were transfected with increasing concentrations (5, 100, 200 nM) of synthetic miR-208b-3p mimics or concentration-matched negative control mimics. While high concentrations of control mimics (particularly 200 nM) induced a non-specific upregulation of both *MYH7* and *MYH6*, miR-208b-3p mimic delivery blunted this non-specific response, maintaining significantly lower *MYH7* levels relative to dose-matched controls (**Figure 3I**). Crucially, this reduction in *MYH7* was unexpected; since miR-208b-3p downregulated known MYH7 repressors, which typically drives *MYH7* expression, *MYH7* levels unexpectedly decreased without the induction of any candidate repressors. Specifically, *PURB*, *CBX1*, *SP3*, and *MED13* exhibited progressive, dose-dependent downregulation following miR-208b-3p mimic treatment, while *SOX6* remained low and *HMOX1* (non-target control) remained unchanged across all conditions (**Figure 3I**). Notably, *ATM*—a predicted target of miR-208b-3p based on a complementary seed-binding site and a mediator of cellular responses to oxidative stress and cytoskeletal mechanics (Bastianello, Porcella et al. 2023, Jiang, Guo et al. 2025)—displayed potent, dose-dependent suppression following miR-208b-3p mimic treatment (**Figure 3I**). Remarkably, *ATM* mRNA was elevated in *Mlp*^⁻/⁻^ myocardium, particularly in females, but showed no comparable increase in human end-stage DCM, with reduced expression observed in ICM (**Supplementary Figure S3E**). This context-dependent expression pattern, together with the suppression observed following miR-208b-3p overexpression, raises the possibility that miR-208b-3p contributes to the regulation of ATM expression during myocardial stress.

Together, these findings argue against a model in which miR-208b-3p regulates *MYH7* solely by depressing its canonical transcriptional repressors. Instead, the targeting of ATM, alongside other previously identified targets involved in stress adaptation, may link miR-208b-3p to the regulation of myocardial redox balance, energy metabolism, and stress-response pathways.

To characterize broader molecular remodeling following miR-208b-3p silencing *in vivo*, transcriptomic profiling and Ingenuity Pathway Analysis (IPA) were performed on myocardium from anti-miR-208b-3p–treated *Mlp^-/-^* mice, scramble-treated *Mlp^-/-^* controls, and wild-type littermates. Suppression of miR-208b-3p led to significant enrichment of pathways regulating redox homeostasis and cell stress-adaptive response, including ERK/MAPK, PI3K/AKT, and sirtuin signaling, alongside activated autophagy (**Figure 3J**). These pathway shifts represented a further elevation of the baseline stress programs observed in untreated *Mlp^-/-^* hearts **(Figure 1B)**. Because autophagy can be induced in response to cellular stress, this heightened activation is consistent with a compensatory response to increased cellular stress following miR-208b-3p suppression.

### miR-208b-3p modulates redox stress and cell survival

To elucidate the mechanism by which miR-208b-3p influences myocardial redox balance, we performed upstream regulator analysis on transcriptomic data from anti-miR-208b-3p–treated *Mlp^-/-^* hearts. This analysis revealed a robust predicted activation of NRF2 (NFE2L2), the master transcriptional regulator of cellular antioxidant defenses (**Figure 4A**). Quantitative PCR confirmed elevation of key NRF2 downstream targets, including *Hmox1* and *Osgin1* (**Figure 4B**). The marked elevation of *Hmox1* indicates an active response to primary oxidative strain, while the elevation of *Osgin1*—an oxidative stress-responsive protein known to drive cytochrome c release and initiate mitochondrial apoptosis (Hu, Yao et al. 2012, Liu, Li et al. 2014)— supports a model in which miR-208b-3p modulates oxidative stress-associated cell-death signaling. Following miR-208b-3p antagomir treatment in *Mlp^-/-^* hearts, malondialdehyde (MDA) levels showed only a modest trend toward increase, while the overall GSH/GSSG ratio remained largely maintained (**Figure 4C–D**). Importantly, *Idh2* expression was not significantly altered following antagomir treatment (**Supplementary Fig. S4A**). Because IDH2 primarily maintains mitochondrial redox balance under chronic oxidative stress (ElBeck, Hossain et al. 2024), this potentially indicates that miR-208b-3p acts through an independent pathway to regulate myocardial redox defenses and pro-apoptotic signaling.

**Figure 4:**
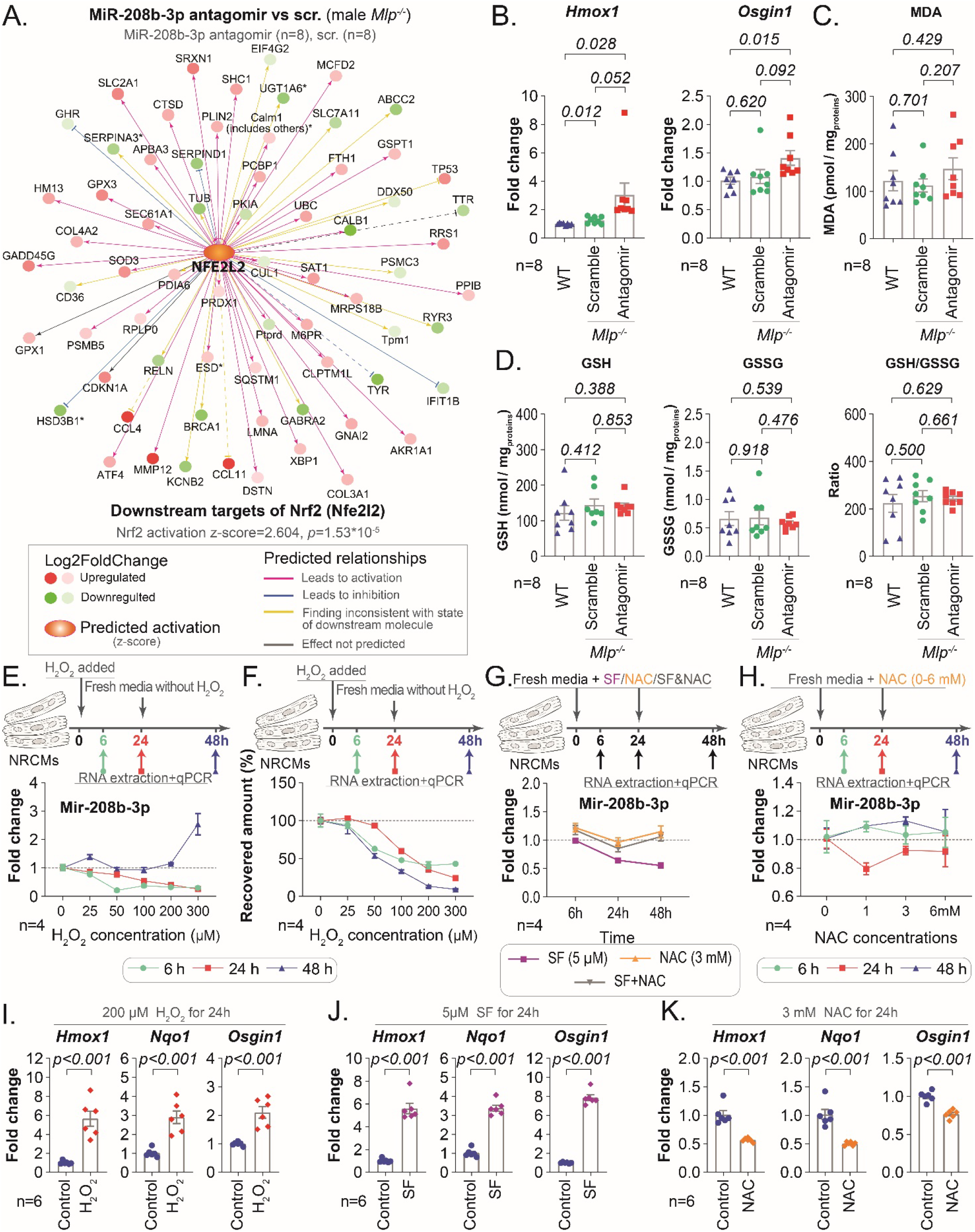
The role of miR-208b-3p in antioxidative response. **A.** IPA upstream regulator analysis of Nrf2 on transcriptomic data from the left ventricle (LV) of *Mlp^-/-^* mice treated with miR-208b-3p antagomirs. **B -D** qPCR analysis of *Hmox1* and *Osgin1* expression and targeted LC-MS/MS analysis of MDA, GSH and GSSG levels in the LV of *Mlp^-/-^* described in figure A. **E.** qPCR analysis of miR-208b-3p expression in NRCMs treated with different concentrations of H2O2 at different time points. **F.** Total RNA was reproduced from (ElBeck, Hossain et al. 2024) and it depicts the recovered amount of extracted RNA from the wells of H2O2-treated NRCMs. **G and H.** qPCR analysis of miR-208b-3p expression in NRCMs treated with different concentrations of sulforaphane (SF) or N-acetylcysteine (NAC) at different time points. **I-K.** qPCR analysis of *Hmox1*, *Nqo1* &*Osgin1* expression in NRCMs treated with 200 µM of H2O2, 5 µM of SF, or 3 mM of NAC, for 24 hours. Bars are the mean values ±SEM with unpaired two-tailed *t*-test.

To test whether endogenous miR-208b-3p directly responds to cellular redox status, primary neonatal rat cardiomyocytes (NRCMs) were exposed to oxidative stimuli *in vitro*. Treatment with increasing doses of H_2_O_2_ (0–300 *μ*M) over 6 and 24 hours triggered a dose-dependent downregulation of miR-208b-3p. Notably, higher concentrations of H_2_O_2_ were associated with substantial cell loss, as reflected by reduced total RNA recovery. Importantly, miR-208b-3p expression was assessed using equal amounts of input RNA, indicating that its downregulation was not simply attributable to differences in the total amount of RNA recovered. Following removal of H_2_O_2_ and incubation in oxidant-free medium for an additional 24 hours, miR-208b-3p expression was restored and exhibited a pronounced rebound induction (**Figure 4E and 4F**).

A similar transient downregulation of miR-208b-3p was observed following treatment with sulforaphane (SF), a milder pro-oxidant electrophile (**Figure 4G**). Co-treatment of SF with the antioxidant N-acetylcysteine (NAC) completely blocked this reduction, restoring miR-208b-3p expression to baseline levels (**Figure 4G**). Furthermore, treatment of NRCMs with NAC alone induced miR-208b-3p expression at lower concentrations (**Figure 4H**). As expected, H_2_O_2_ and SF strongly activated NRF2 downstream target genes, whereas NAC suppressed them (**Figure 4I–K**).

The acute suppression of miR-208b-3p under lethal oxidative challenge, coupled with its rapid rebound in surviving cells upon oxidant removal or exposure to reducing agents, indicates that miR-208b-3p dynamically regulated in response to changes in intracellular redox state. Together with its modulation of pro-apoptotic factors like *Osgin1*, these findings support a model in which miR-208b-3p acts as a temporal survival brake during severe oxidative strain, restraining pro-apoptotic signaling to provide cardiomyocytes a critical window to clear damaged organelles, activate NRF2-mediated defenses, and restore redox homeostasis.

### Loss of miR-208b-3p activates multiple oxidative stress–responsive signaling pathways independent of direct NRF2 feedback

Although knockdown of miR-208b-3p in *Mlp^-/-^* myocardium activated NRF2 (**Figure 4A**), we next tested whether direct NRF2 activation feedback-regulates miR-208b-3p expression. Pharmacological activation of NRF2 in *Mlp^-/-^* hearts via AZ925—a small molecule that disrupts the KEAP1–NRF2 interaction to improve cardiac function in male *Mlp^-/-^* mice (ElBeck, Hossain et al. 2024)—did not alter myocardial miR-208b-3p expression (**Figure 5A-D**). These findings demonstrate that NRF2 is not a direct upstream transcriptional regulator of miR-208b-3p. Instead, miR-208b-3p and NRF2 may operate as parallel, interconnected arms of the cardiac oxidative stress response.

**Figure 5:**
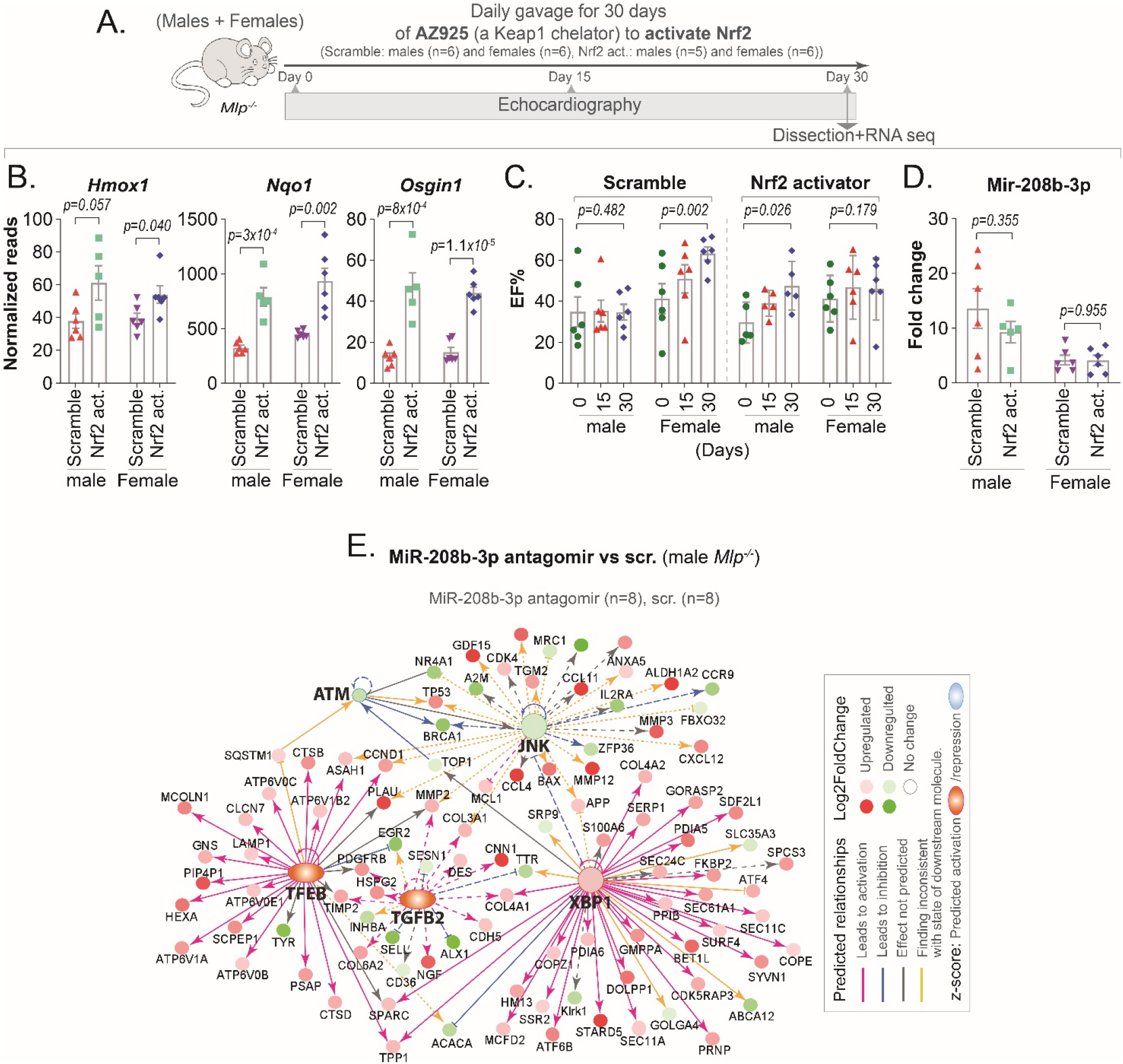
miR208b-3p activates adaptive stress response. **A.** Schematic illustration of the *in vivo* Keap1 chelator (Nrf2 activator) study design. **B-D.** Expression levels of NRF2 downstream targets *Hmox1*, *Nqo1*, and *Osgin1* estimated by RNA seq in (**B**), ejection fraction (EF%) estimated by echocardiography in (**C**), and qPCR analysis of miR-208b-3p expression in (**D**), in the LV of *Mlp^-/-^* treated with Nrf2 activator, normalized to scramble control treated animals. **E.** IPA *in silico* upstream regulator analysis of top stress-responsive regulators on transcriptomic data from the myocardium of *Mlp^-/-^* mice treated with miR-208b-3p antagomirs. Bars are the mean values ±SEM with unpaired two-tailed *t*-test in figures **B** and **D**, and with paired one-tailed *t*-test in figure **C**. Figures **B** and **C** are adapted from (ElBeck, Hossain et al. 2024).

To further characterize the transcriptional network responsive to miR-208b-3p inhibition, we performed IPA upstream regulator analysis on RNA-sequencing data from miR-208b-3p antagomir-treated *Mlp^-/-^*hearts. This analysis predicted robust activation of multiple key stress- and damage-responsive regulators, including TFEB (master regulator of lysosomal biogenesis and autophagy), XBP1 (endoplasmic reticulum stress sensor), TGFB1, and JNK signaling, alongside signature networks linked to reactive oxygen species and hydrogen peroxide accumulation (**Figure 5E**).

### miR-208b-3p modulates oxidative stress–associated apoptosis and autophagy *in vitro*

Given that miR-208b-3p knockdown elevated the pro-apoptotic factor *Osgin1* (**Figure 4B**) and induced mild predicted activation of p53 target signaling (**Figure 3J**), we sought to determine whether miR-208b-3p directly restrains pro-apoptotic cascades under stress. Mapping transcriptomic data from *Mlp^-/-^* hearts onto the canonical Apoptosis Signaling pathway revealed widespread baseline inhibition of downstream apoptotic nodes in *Mlp^-/-^* myocardium, whereas *in vivo* knockdown of miR-208b-3p reversed this signature, predicting reactivation of multiple pro-apoptotic signaling hubs (**Supplementary Figure S6A**).

To further examine these findings in a controlled setting, miR-208b-3p was overexpressed in HEK293 cells, followed by RNA sequencing and transcriptomic pathway analysis. Differentially expressed genes mapped onto a curated NRF2/oxidative stress response network and were associated with predicted suppression of oxidative stress-related apoptotic signaling (**Figure 6A**). In parallel, network analysis predicted enrichment of autophagy pathways, consistent with activation of cytoprotective stress-response programs. To determine whether previously reported miR-208b-3p target genes are functionally connected to this redox-defense network, established target genes were overlaid onto the interaction network. Several reported targets—including *SOX6*, *PURB*, *CBX1*, *SP3*, and *ETS1* (van Rooij, Quiat et al. 2009, Itoh, Takeda et al. 2010)—showed extensive functional predicted interactions with NRF2 and oxidative stress-associated network nodes (**Figure 6A**). These findings suggest that miR-208b-3p targets intersect with a broader cytoprotective network involving redox regulation, autophagy, and oxidative stress–associated apoptosis. In particular, *ETS1* (Xu, Wilson et al. 2002), together with *SOX6*, *PURB*, *CBX1*, and *SP3*, may contribute to modulation of oxidative stress–associated apoptotic signaling, with *ATM* providing additional network-level support for their integration into these pathways.

**Figure 6:**
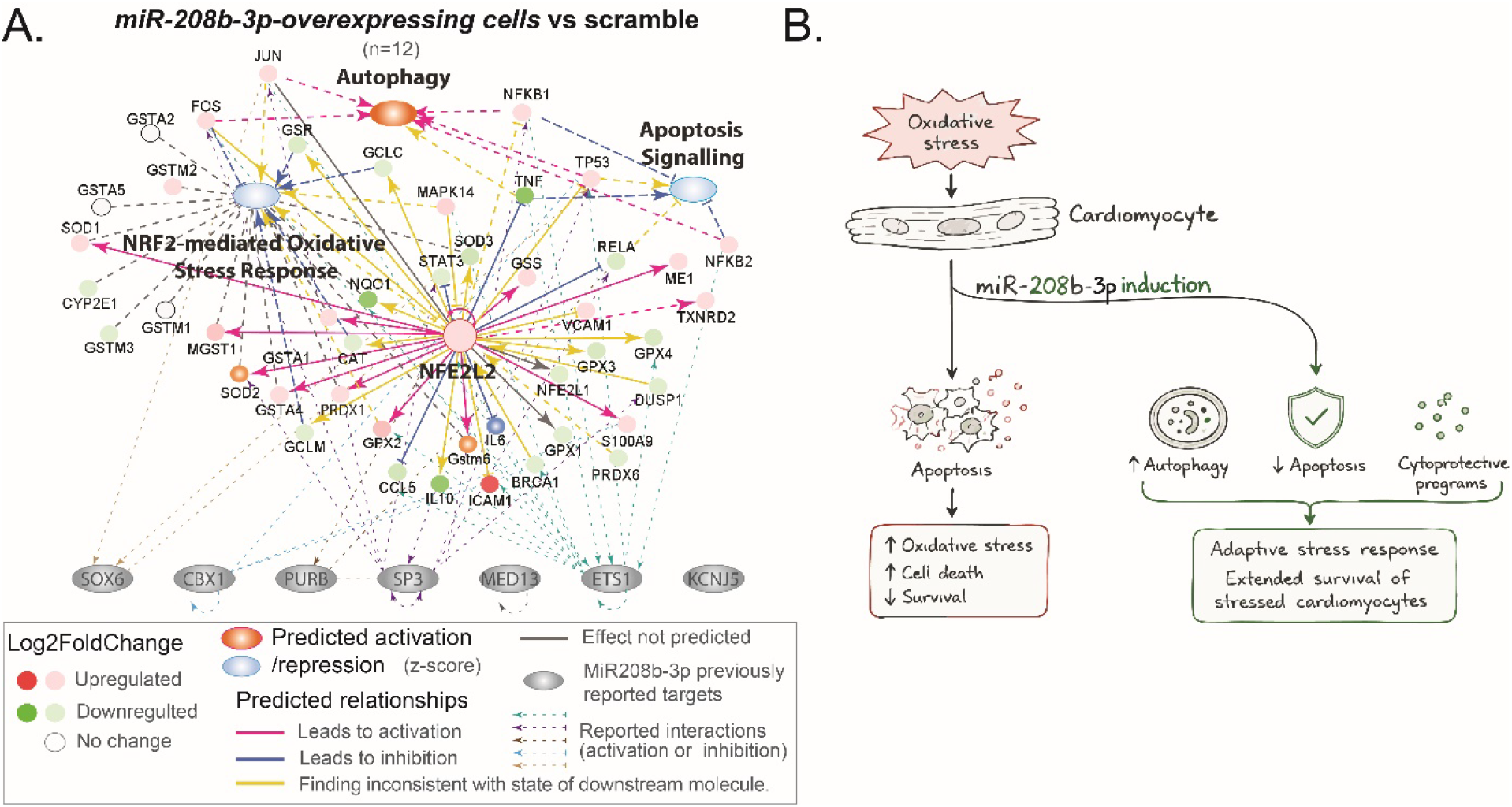
miR208b-3p activates adaptive stress response and attenuates apoptosis signaling. **A.** IPA in silico interaction network linking differentially expressed genes from miR-208b-3p-overexpressing HEK293 cells with oxidative stress-associated apoptosis and autophagy through the NRF2-mediated oxidative stress response. Interactions with previously identified targets of miR208b-3p were converted to dotted lines to simplify the figure. **B.** Schematic summary of the proposed role of miR-208b-3p in the cardiomyocyte stress-adaptation response.

Ultimately, these findings demonstrate that miR-208b-3p functions as an adaptive, stress-induced brake on oxidative stress associated programmed cell death. By dampening primary pro-apoptotic nodes, upstream oxidative stress, and stress-sensing pathways, miR-208b-3p prevents the cellular damage that forces secondary stress-response surges, providing cardiomyocytes with a critical temporal window to manage bioenergetic strain, restore redox homeostasis, and attempt structural repair (**Figure 6B**).

## Discussion

Our findings establish miR-208b-3p as an upstream regulator of redox-associated cardiomyocyte survival and responses in the failing heart. By studying genetic DCM models characterized by stress adapted—exemplified by *Mlp^-/-^* mice with their marked survival and pronounced miR-208b-3p induction—we found that silencing miR-208b-3p *in vivo* represses pro-survival networks and unleashes oxidative-response genes such as *Osgin1*. Conversely, endogenous miR-208b-3p becomes preferentially elevated in cardiomyocytes that survived extreme pro-oxidant exposure. These dynamics hold significant translational relevance for clinical myocardial infarction, where circulating miR-208b-3p spikes alongside myocardial oxidative stress (Grieve, Byrne et al. 2004, Hori and Nishida 2009, Corsten, Dennert et al. 2010, Gidlof, Andersson et al. 2011). The initial downregulation of miR-208b-3p during acute oxidative insult allows for the immediate mounting of NRF2-mediated antioxidant defenses. However, its subsequent rebound as oxidative stress subsides acts as a functional feedback loop: by fine-tuning late-stage antioxidant signaling, miR-208b-3p prevents the hyper-activation of stress pathways that would otherwise trigger apoptosis. The modest induction of miR-208b-3p by reducing agents such as N-acetylcysteine further confirms its sensitivity to intracellular redox state.

While miR-208b-3p inhibition in *Mlp*^⁻/⁻^ hearts was associated with increased expression of NRF2 target genes, pharmacological NRF2 activation did not alter miR-208b-3p expression, suggesting that this pathway may represent a parallel or upstream component of the myocardial stress response. miR-208b-3p overexpression also affected several genes implicated in stress adaptation and cell survival, including ATM, ETS1 and other previous identified targets of miR-208b-3p. ATM has been linked to oxidative stress responses, NRF2 signaling, and cytoskeletal regulation (Bastianello, Porcella et al. 2023, Lee 2024, Jiang, Guo et al. 2025), but its effects in cardiac injury appear context-dependent. Although ATM suppression has been associated with reduced apoptosis and fibrosis-related signaling in experimental myocardial infarction (Chiang, Liang et al. 2020), ATM deficiency can also impair cardiac recovery (Foster, Daniel et al. 2013, Daniel, Daniels et al. 2014, Daniel, Scofield et al. 2016). In our study, miR-208b-3p overexpression suppressed *ATM*, whose expression was elevated in *Mlp*⁻/⁻ myocardium, particularly in females, which exhibit enhanced stress adaptation compared with males (ElBeck, Hossain et al. 2024). In human end-stage heart failure, *ATM* expression remained unchanged in DCM but was reduced in ICM, which is generally associated with a poorer prognosis and higher mortality than DCM. Given the context-dependent role of ATM in cardiac injury, these findings suggest that miR-208b-3p modulates the abundance of ATM alongside other stress-related targets, including ETS1, within a tightly regulated stress-response network to support redox balance and cell survival during myocardial remodeling. Further studies are needed to establish the contribution of these targets to the effects of miR-208b-3p.

The observation that miR-208b-3p knockdown reduces *Myh7* transcript levels without derepressing canonical transcriptional repressors points to an indirect mechanism of myosin isoform regulation that may be linked to cellular metabolic and stress-response pathways. Rather than operating through a direct repressor network, *Myh7* induction in the failing heart serves as a metabolic adaptation; *β*-myosin heavy chain utilizes ATP more efficiently than *α*-myosin under conditions of energetic and oxidative strain (Hasenfuss, Mulieri et al. 1991, Rassier and Mansson 2025). By targeting upstream oxidative nodes such as ATM, miR-208b-3p serves as a primary upstream checkpoint against intracellular redox stress. Suppressing miR-208b-3p disrupts this protective shield, modestly increasing primary lipid peroxidation and triggering a secondary, reactive activation of sirtuin, ERK/MAPK, and autophagy networks to process the resulting organellar damage. Consequently, the concurrent downregulation of *Myh7* following miR-208b-3p suppression does not reflect direct transcriptional repatterning, but rather a secondary shifting of the cardiac stress program as the myocardium engages compensatory clearance mechanisms to manage the heightened oxidative strain.

In conclusion, our findings identify miR-208b-3p as a regulator of redox-associated stress responses and apoptosis-associated signaling in the remodeling myocardium. The contrasting activation of EIF2 signaling, autophagy, and the unfolded protein response in *Mlp*^⁻/⁻^ hearts versus their attenuation in human end-stage DCM highlights a fundamental difference between stress-adapted myocardium and advanced human heart failure. Together, these results establish miR-208b-3p as a candidate mediator of cardiac stress adaptation and provide a basis for further investigating its role in cardiomyocyte survival and myocardial remodeling.

## Material and methods

### Study Design and Experimental Overview

This study used complementary *in vivo*, *in vitro*, and transcriptomic approaches to investigate the role of miR-208b-3p in myocardial stress adaptation and remodeling. Myocardial miR-208b-3p expression and transcriptomic profiles were examined in *Mlp^−/−^* mice and human DCM and ICM myocardium. The functional effects of miR-208b-3p inhibition were assessed in *Mlp^−/−^* mice treated with an LNA-modified miR-208b-3p antagomir, with cardiac function and myocardial molecular responses evaluated longitudinally. Complementary *in vitro* gain-of-function experiments were performed, and the miR-208b-3p responses to oxidative and reducing conditions were examine. Transcriptomic, pathway, and biochemical analyses were used to characterize the molecular responses associated with miR-208b-3p modulation.

Detailed experimental procedures are provided in the accompanying Materials and Methods document.

## Supporting information

Detailed Material and Methods

## Acknowledgments

We thank Professor Christer Betsholtz, Professor Mattias Svensson, Dr. Akos Vegvari, Dr. Eike-Benjamin Braune and Dr. Mukesh Varshney at Karolinska Institutet for their support and valuable contributions. We also thank Dr. Juliana Nardelli and Dr. Karin Sundström at QIAGEN for their support and guidance for the experimental design of the studies involving the miR-208b-3p antagomirs and mimics. Finally, we acknowledge the Single Cell Core Facility in Flemingsberg for their assistance and technical support. The authors would like to acknowledge the contributions of the Small Molecule Mass Spectrometry Core Facility (KI-SMMS), financed by the Infrastructure Board at Karolinska Institutet for providing scientific input and analysis of the samples. We would like to acknowledge the Single-cell core facility at the Flemingsberg campus (SICOF) at Karolinska Institutet (KI) for their sequencing. Portions of this manuscript and data were adapted from the author’s PhD thesis, [Redox and epigenetic modulators regulate cardiac function and remodeling in health and disease], Karolinska Institutet, 2024. Generative AI was used to assist with the language of the manuscript. The authors reviewed and edited the resulting text and take full responsibility for the final content.

## Contributions

M.B.H. performed echocardiography, contributed to the design of the in vivo studies, conducted the IPA analyses, provided study materials, interpreted the findings, and revised the manuscript. H.S. performed echocardiography and bioinformatic analyses. Z.E. conceived and designed the study, carried out and supervised the experiments, led the data analysis and interpretation, and drafted the manuscript.

## References

Akat, K. M., D. Moore-McGriff, P. Morozov, M. Brown, T. Gogakos, J. Correa Da Rosa, A. Mihailovic, M. Sauer, R. Ji, A. Ramarathnam, H. Totary-Jain, Z. Williams, T. Tuschl and P. C. Schulze (2014). "Comparative RNA-sequencing analysis of myocardial and circulating small RNAs in human heart failure and their utility as biomarkers." Proc Natl Acad Sci U S A 111(30): 11151–11156.

Ambardekar, A. V. and P. M. Buttrick (2011). "Reverse remodeling with left ventricular assist devices: a review of clinical, cellular, and molecular effects." Circ Heart Fail 4(2): 224–233.

Arber, S., J. J. Hunter, J. Ross, Jr., M. Hongo, G. Sansig, J. Borg, J. C. Perriard, K. R. Chien and P. Caroni (1997). "MLP-deficient mice exhibit a disruption of cardiac cytoarchitectural organization, dilated cardiomyopathy, and heart failure." Cell 88(3): 393–403.

Aurora, A. B., A. I. Mahmoud, X. Luo, B. A. Johnson, E. van Rooij, S. Matsuzaki, K. M. Humphries, J. A. Hill, R. Bassel-Duby, H. A. Sadek and E. N. Olson (2012). "MicroRNA-214 protects the mouse heart from ischemic injury by controlling Ca(2)(+) overload and cell death." J Clin Invest 122(4): 1222–1232.

Bastianello, G., G. Porcella, G. V. Beznoussenko, G. Kidiyoor, F. Ascione, Q. Li, A. Cattaneo, V. Matafora, A. Disanza, M. Quarto, A. A. Mironov, A. Oldani, S. Barozzi, A. Bachi, V. Costanzo, G. Scita and M. Foiani (2023). "Cell stretching activates an ATM mechano-transduction pathway that remodels cytoskeleton and chromatin." Cell Rep 42(12): 113555.

Burkhoff, D., V. K. Topkara, G. Sayer and N. Uriel (2021). "Reverse Remodeling With Left Ventricular Assist Devices." Circ Res 128(10): 1594–1612.

Carbonell, T. and A. V. Gomes (2020). "MicroRNAs in the regulation of cellular redox status and its implications in myocardial ischemia-reperfusion injury." Redox Biol 36: 101607.

Cheng, X., C. H. Ku and R. C. Siow (2013). "Regulation of the Nrf2 antioxidant pathway by microRNAs: New players in micromanaging redox homeostasis." Free Radic Biol Med 64: 4–11.

Chiang, M. H., C. J. Liang, L. C. Lin, Y. F. Yang, C. C. Huang, Y. H. Chen, H. L. Kao, Y. C. Chen, S. R. Ke, C. W. Lee, M. S. Lin and Y. L. Chen (2020). "miR-26a attenuates cardiac apoptosis and fibrosis by targeting ataxia-telangiectasia mutated in myocardial infarction." J Cell Physiol 235(9): 6085–6102.

Corsten, M. F., R. Dennert, S. Jochems, T. Kuznetsova, Y. Devaux, L. Hofstra, D. R. Wagner, J. A. Staessen, S. Heymans and B. Schroen (2010). "Circulating MicroRNA-208b and MicroRNA-499 reflect myocardial damage in cardiovascular disease." Circ Cardiovasc Genet 3(6): 499–506.

Daniel, L. L., C. R. Daniels, S. Harirforoosh, C. R. Foster, M. Singh and K. Singh (2014). "Deficiency of ataxia telangiectasia mutated kinase delays inflammatory response in the heart following myocardial infarction." J Am Heart Assoc 3(6): e001286.

Daniel, L. L., S. L. Scofield, P. Thrasher, S. Dalal, C. R. Daniels, C. R. Foster, M. Singh and K. Singh (2016). "Ataxia telangiectasia-mutated kinase deficiency exacerbates left ventricular dysfunction and remodeling late after myocardial infarction." Am J Physiol Heart Circ Physiol 311(2): H445–452.

ElBeck, Z., M. B. Hossain, H. Siga, N. Oskolkov, F. Karlsson, J. Lindgren, A. Walentinsson, D. Koppenhofer, R. Jarvis, R. Burli, T. Jamier, E. Franssen, M. Firth, A. Degasperi, C. Bendtsen, R. I. Menzies, K. Streckfuss-Bomeke, M. Kohlhaas, A. G. Nickel, L. H. Lund, C. Maack, A. Vegvari and C. Betsholtz (2024). "Epigenetic modulators link mitochondrial redox homeostasis to cardiac function in a sex-dependent manner." Nat Commun 15(1): 2358.

Fierro-Fernandez, M., V. Miguel and S. Lamas (2016). "Role of redoximiRs in fibrogenesis." Redox Biol 7: 58–67.

Foster, C. R., L. L. Daniel, C. R. Daniels, S. Dalal, M. Singh and K. Singh (2013). "Deficiency of ataxia telangiectasia mutated kinase modulates cardiac remodeling following myocardial infarction: involvement in fibrosis and apoptosis." PLoS One 8(12): e83513.

Gidlof, O., P. Andersson, J. van der Pals, M. Gotberg and D. Erlinge (2011). "Cardiospecific microRNA plasma levels correlate with troponin and cardiac function in patients with ST elevation myocardial infarction, are selectively dependent on renal elimination, and can be detected in urine samples." Cardiology 118(4): 217–226.

Grieve, D. J., J. A. Byrne, A. C. Cave and A. M. Shah (2004). "Role of oxidative stress in cardiac remodelling after myocardial infarction." Heart Lung Circ 13(2): 132–138.

Hasenfuss, G., L. A. Mulieri, E. M. Blanchard, C. Holubarsch, B. J. Leavitt, F. Ittleman and N. R. Alpert (1991). "Energetics of isometric force development in control and volume-overload human myocardium. Comparison with animal species." Circ Res 68(3): 836–846.

Hori, M. and K. Nishida (2009). "Oxidative stress and left ventricular remodelling after myocardial infarction." Cardiovasc Res 81(3): 457–464.

Hu, J., H. Yao, F. Gan, A. Tokarski and Y. Wang (2012). "Interaction of OKL38 and p53 in regulating mitochondrial structure and function." PLoS One 7(8): e43362.

Itoh, T., S. Takeda and Y. Akao (2010). "MicroRNA-208 modulates BMP-2-stimulated mouse preosteoblast differentiation by directly targeting V-ets erythroblastosis virus E26 oncogene homolog 1." J Biol Chem 285(36): 27745–27752.

Jiang, X. Y., Q. Q. Guo, S. S. Wang, R. Guo, Y. Zou, J. W. Liu, Y. L. Feng, Y. Guo, Y. H. Li, X. Y. Liu, X. Y. Zhang, S. Hao, X. X. Wu, M. H. Li, A. Liu, C. L. Li, W. D. Guo, H. D. Xu, X. Y. Song, T. Finkel and L. Cao (2025). "DNA damage response pathway regulates Nrf2 in response to oxidative stress." Sci Adv 11(30): eadu9555.

Kurinna, S. and S. Werner (2015). "NRF2 and microRNAs: new but awaited relations." Biochem Soc Trans 43(4): 595–601.

Lee, J. H. (2024). "Oxidative stress and the multifaceted roles of ATM in maintaining cellular redox homeostasis." Redox Biol 75: 103269.

Liu, M., Y. Li, L. Chen, T. H. Chan, Y. Song, L. Fu, T. T. Zeng, Y. D. Dai, Y. H. Zhu, Y. Li, J. Chen, Y. F. Yuan and X. Y. Guan (2014). "Allele-specific imbalance of oxidative stress-induced growth inhibitor 1 associates with progression of hepatocellular carcinoma." Gastroenterology 146(4): 1084–1096.

Manea, S. A., A. Constantin, G. Manda, S. Sasson and A. Manea (2015). "Regulation of Nox enzymes expression in vascular pathophysiology: Focusing on transcription factors and epigenetic mechanisms." Redox Biol 5: 358–366.

Mathiyalagan, P., J. Okabe, L. Chang, Y. Su, X. J. Du and A. El-Osta (2014). "The primary microRNA-208b interacts with Polycomb-group protein, Ezh2, to regulate gene expression in the heart." Nucleic Acids Res 42(2): 790–803.

Palmer, B. M. (2005). "Thick filament proteins and performance in human heart failure." Heart Fail Rev 10(3): 187–197.

Rassier, D. E. and A. Mansson (2025). "Mechanisms of myosin II force generation: insights from novel experimental techniques and approaches." Physiol Rev 105(1): 1–93.

Satoh, M., Y. Minami, Y. Takahashi, T. Tabuchi and M. Nakamura (2010). "Expression of microRNA-208 is associated with adverse clinical outcomes in human dilated cardiomyopathy." J Card Fail 16(5): 404–410.

Selvam, M., V. Bandi, S. Ponne, C. Ashok and S. Baluchamy (2022). "microRNA-150 targets major epigenetic repressors and inhibits cell proliferation." Exp Cell Res 415(1): 113110.

Sielemann, K., Z. Elbeck, A. Gartner, A. Brodehl, C. Stanasiuk, H. Fox, L. Paluszkiewicz, J. Tiesmeier, S. Wlost, J. Gummert, S. P. Albaum, J. Sielemann, R. Knoll and H. Milting (2020). "Distinct Myocardial Transcriptomic Profiles of Cardiomyopathies Stratified by the Mutant Genes." Genes (Basel) 11(12).

Sozen, E., B. Karademir and N. K. Ozer (2015). "Basic mechanisms in endoplasmic reticulum stress and relation to cardiovascular diseases." Free Radic Biol Med 78: 30–41.

Sucharov, C. C., D. P. Kao, J. D. Port, A. Karimpour-Fard, R. A. Quaife, W. Minobe, K. Nunley, B. D. Lowes, E. M. Gilbert and M. R. Bristow (2017). "Myocardial microRNAs associated with reverse remodeling in human heart failure." JCI Insight 2(2): e89169.

Sweet, M. E., A. Cocciolo, D. Slavov, K. L. Jones, J. R. Sweet, S. L. Graw, T. B. Reece, A. V. Ambardekar, M. R. Bristow, L. Mestroni and M. R. G. Taylor (2018). "Transcriptome analysis of human heart failure reveals dysregulated cell adhesion in dilated cardiomyopathy and activated immune pathways in ischemic heart failure." BMC Genomics 19(1): 812.

van Heesch, S., F. Witte, V. Schneider-Lunitz, J. F. Schulz, E. Adami, A. B. Faber, M. Kirchner, H. Maatz, S. Blachut, C. L. Sandmann, M. Kanda, C. L. Worth, S. Schafer, L. Calviello, R. Merriott, G. Patone, O. Hummel, E. Wyler, B. Obermayer, M. B. Mucke, E. L. Lindberg, F. Trnka, S. Memczak, M. Schilling, L. E. Felkin, P. J. R. Barton, N. M. Quaife, K. Vanezis, S. Diecke, M. Mukai, N. Mah, S. J. Oh, A. Kurtz, C. Schramm, D. Schwinge, M. Sebode, M. Harakalova, F. W. Asselbergs, A. Vink, R. A. de Weger, S. Viswanathan, A. A. Widjaja, A. Gartner-Rommel, H. Milting, C. Dos Remedios, C. Knosalla, P. Mertins, M. Landthaler, M. Vingron, W. A. Linke, J. G. Seidman, C. E. Seidman, N. Rajewsky, U. Ohler, S. A. Cook and N. Hubner (2019). "The Translational Landscape of the Human Heart." Cell 178(1): 242–260 e229.

van Rooij, E., D. Quiat, B. A. Johnson, L. B. Sutherland, X. Qi, J. A. Richardson, R. J. Kelm, Jr. and E. N. Olson (2009). "A family of microRNAs encoded by myosin genes governs myosin expression and muscle performance." Dev Cell 17(5): 662–673.

Xu, D., T. J. Wilson, D. Chan, E. De Luca, J. Zhou, P. J. Hertzog and I. Kola (2002). "Ets1 is required for p53 transcriptional activity in UV-induced apoptosis in embryonic stem cells." EMBO J 21(15): 4081–4093.

