## Supplementary figures for "MiR-208b-3p is an upstream regulator of redox-associated cardiomyocyte survival"

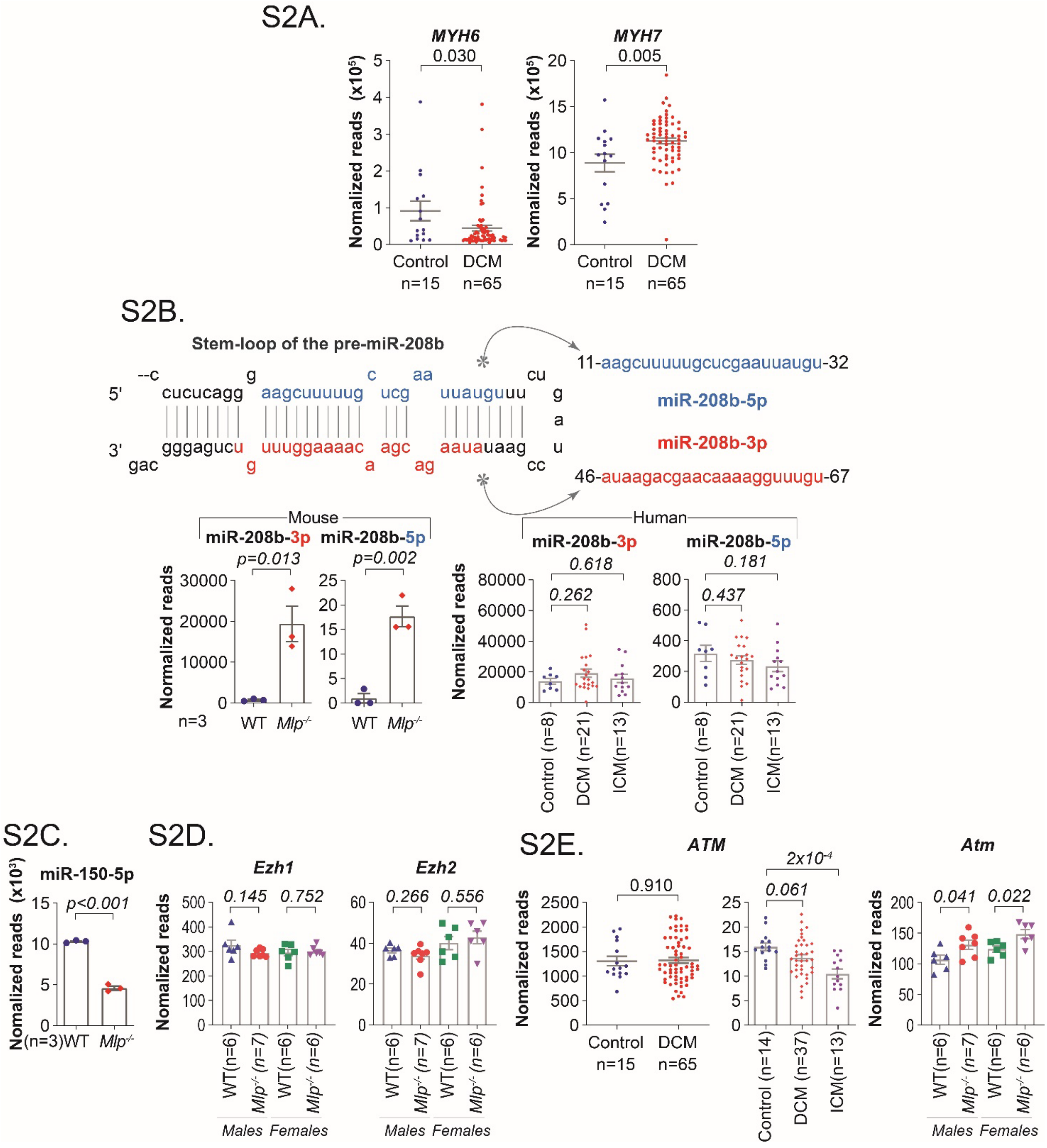


**Supplementary Figure 2. Additional analysis of miRNA expression in *Mlp*$^{-/-}$ myocardium and human cardiomyopathy.**

**A.** *MYH6* and *MYH7* expression in the LV of patients with eccentric DCM extracted from a public repository (van Heesch, Witte et al. 2019). **B.** Stem loop of pre-miR-208b, and the expression levels of its 5p and 3p mature miRNAs in WT and *Mlp*$^{-/-}$ myocardium and in human control, DCM, and ICM myocardium estimated by small RNA seq described in main figure 2. Human small RNA-seq data were adapted from Akat et al. (Akat, Moore-McGriff et al. 2014).The plot for miR-208b-3p in human control, DCM, and ICM myocardium was replicated from main figure 2C to enable an easy comparison of the intensity of expression with miR-208b-3p here. **C.** miR-150-5p expression in WT and *Mlp*$^{-/-}$ myocardium by small RNA-seq. **D.** Expression levels of *Ezh1* and *Ezh2* in the LV of WT and *Mlp*$^{-/-}$ extracted from a public repository (ElBeck, Hossain et al. 2024). **E.** *ATM* expression in the LV of patients with eccentric DCM and in the LV of WT and *Mlp*$^{-/-}$ extracted from a public repository (Sweet, Cocciolo et al. 2018, van Heesch, Witte et al. 2019, ElBeck, Hossain et al. 2024) Data are mean ± SEM; individual points represent individual samples. Statistical analysis was performed using unpaired two-tailed *t*-tests.

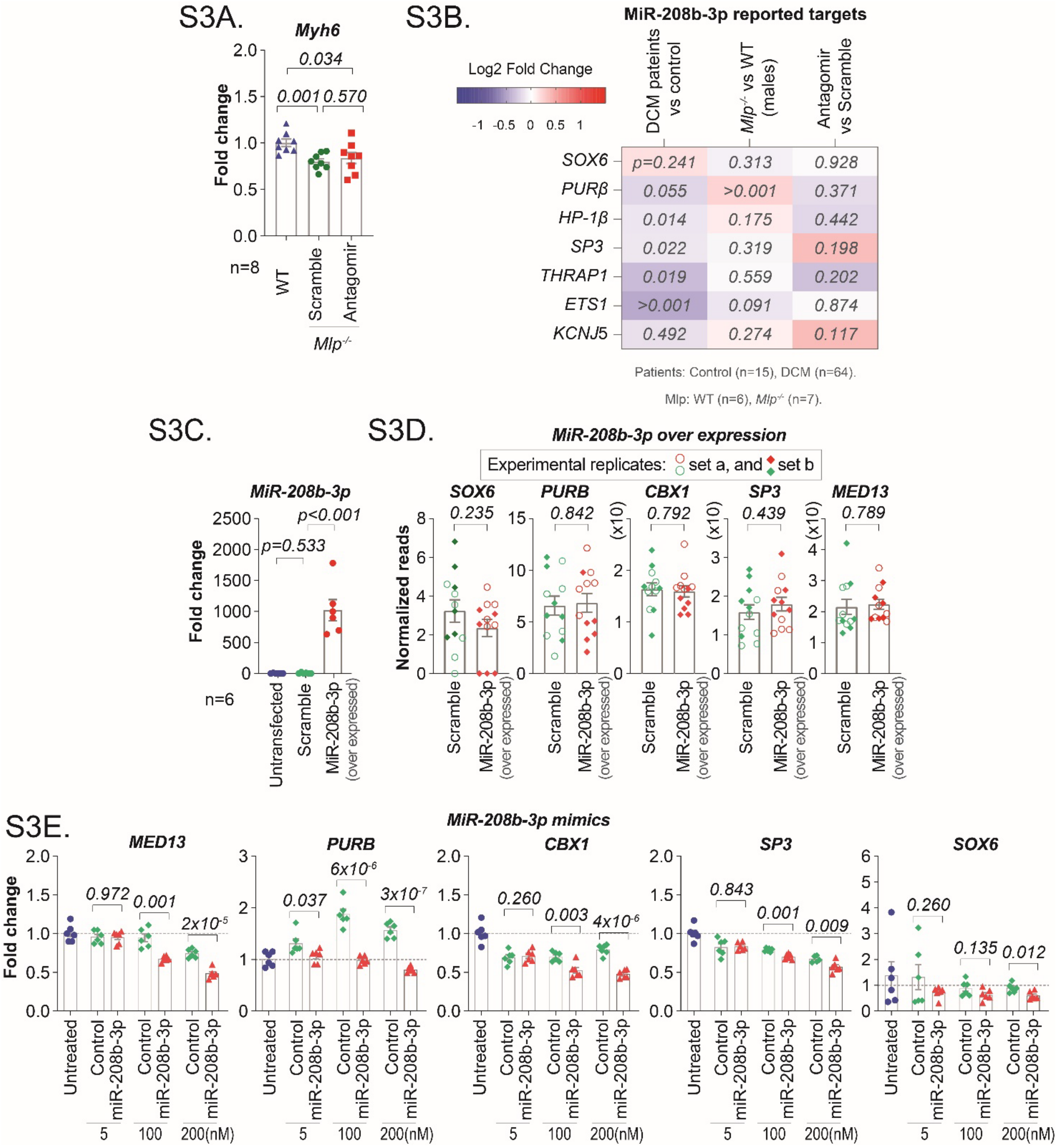


**Supplementary Figure 3. Molecular effects of miR-208b-3p inhibition and induction.**

**A.** *Myh6* expression in LV myocardium following miR-208b-3p antagomir treatment, normalized to untreated WT littermates. **B.** Expression of previously reported targets of miR-208b-3p in the myocardium of *Mlp*$^{-/-}$ mice treated with miR-208b-3p antagomirs, as well as, in patients with DCM and untreated *Mlp*$^{-/-}$ assessed by RNA sequencing. Transcriptomic data of patients and untreated *Mlp*$^{-/-}$ were obtained from public sources (van Heesch, Witte et al. 2019) and (ElBeck, Hossain et al. 2024), respectively. Numbers in *italic* inside the boxes indicate *p*-values. **C.** qPCR analysis of miR-208b-3p expression in HEK293 cells upon transfecting them with mammalian shRNA vector bearing a miR-208b-3p Hairpin Sequence, or a scramble vector expressing EGFP. **D.** Expression levels of some of the previously reported miR-208b-3p targets and their downstream target in HEK293 cells overexpressing miR-208b-3p assessed by RNA sequencing. **E.** *MED13, PURB, CBX1, SP3* and *SOX6* expression in HEK293 cells treated with 5, 100, or 200 nM miR-208b-3p mimic or concentration-matched negative control mimic. Data are mean ± SEM; individual points represent individual samples. Statistical analysis was performed using unpaired two-tailed *t*-tests.

S4A.

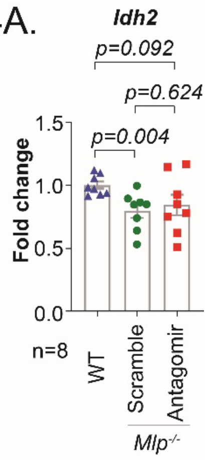


**Supplementary Figure 4. Idh2 expression following miR-208b-3p inhibition in *Mlp*$^{-/-}$ myocardium**

**A** qPCR analysis of *Idh2* in the left ventricle (LV) of *Mlp*$^{-/-}$ mice treated with miR-208b-3p antagomirs.

S6A.

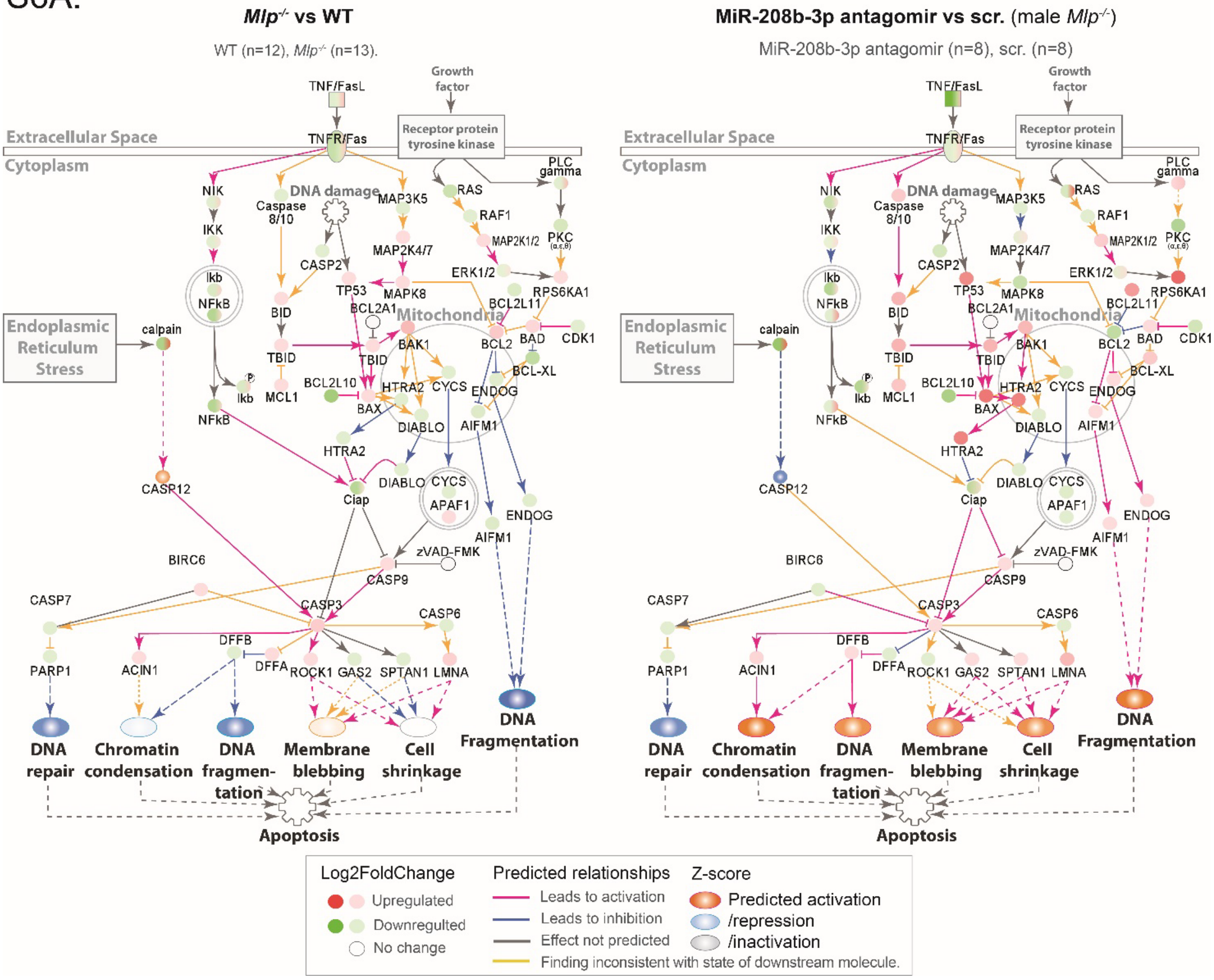


**Supplementary figure S6: knockdown of miR-208b-3p activates apoptosis signalling**

**S6A.** IPA Apoptosis Signaling canonical pathway overlaid with DEGs from transcriptomic data from the myocardium of *Mlp*$^{-/-}$ mice compared to WT controls and from the myocardium of *Mlp*$^{-/-}$ mice treated with miR-208b-3p antagomirs in comparison to scramble treated controls. Transcriptomic data of untreated *Mlp*$^{-/-}$ were obtained from public (ElBeck, Hossain et al. 2024).
