## Supplementary material for "MiR-208b-3p is an upstream regulator of redox-associated cardiomyocyte survival": Detailed Material and Methods

**Detailed martial and methods utilized in**

### 1. Animals

*Mlp*$^{-/-}$ mice, also known as *Csrp3*$^{-/-}$ mice, were maintained on a hybrid 129/Sv and C57BL/6N genetic background and bred in-house. Heterozygous animals were intercrossed to generate wild-type (WT) and *Mlp*$^{-/-}$ littermates. Male *Mlp*$^{-/-}$ mice were used for the miR-208b-3p antagomir studies and the corresponding molecular analyses. Mice were housed under a 12-hour light/dark cycle with free access to standard chow and water at an ambient temperature of 20–24°C and 45–65% humidity. Primary neonatal rat cardiomyocytes (NRCMs) were isolated from hearts of 3-day-old Sprague-Dawley rats. All animal experiments were conducted in accordance with European Union legislation governing the protection of animals used for scientific purposes (Directive 2010/63/EU) and were approved by the local animal ethics committee in Linköping, Sweden (permit numbers 1369 and 2713).

#### 1.1. *In vivo* miR-208b-3p antagomir study

LNA-modified antagomir against miR-208b-3p and a scrambled control sequence were obtained from Qiagen (miRCURY LNA™) (Table 1). The study was designed and performed in accordance with the PREPARE guidelines (Smith, Clutton et al. 2018).

**Table 1: The sequence of utilized antagomirs and scramble control.**

| Target | sequences | Cat. No |
|---|---|---|
| I-MMU-MIR-208B-3P | C*T*T*T*T*G*T*T*C*G*T*C*T*T*A | 339203 YCI0201031-FZA |
| MMU-MIR-SCRAMBLE | A*C*G*T*C*T*A*T*A*C*G*C*C*C*A | 339203 YCI0202044-FZA |

* indicate phosphorothioate bonds

The predicted improvement of cardiac function in *Mlp*$^{-/-}$ mice was small, due to the extensive variability in the disease severity of these animals (ElBeck, Hossain et al. 2024). We therefore argued that useful data could not be achieved from treating just a few animals. Hence, we skipped the pilot step, and instead directly treated 8 animals in each group (antagomir/scramble), which we estimated to be enough to observe a significant improvement in the cardiac function if lowering miR-208b-3p level would have beneficial effects. This assumption was based on our previous experience from similar experiments.

*Mlp*$^{-/-}$ animals, at the age of 8 weeks, were randomized into two groups based on their body weights (see Figure 3A for the experimental design of the study). Each animal received 4 injections of 80 mg/kg of antagomir or scrambled control at days 0, 7, 14 and 21. Echocardiography was performed on days 0, 17 and 28. The study was terminated immediately after the last echocardiographic examination, and the hearts were dissected and frozen immediately for subsequent biochemical analyses. No animals died before terminating the study, and there were no observable changes on their behaviors or body weight. The experimenters were blinded to the treatment groups during different procedures. The examination with echocardiography was performed with the exact same procedure described in Elbeck *et al.* 2024 (ElBeck, Hossain et al. 2024). Briefly, mice were anesthetized with 3% isoflurane and maintained at 1.5% isoflurane during imaging. Echocardiography was performed using a Vevo 3100 ultrasound system with an MX550D probe (VisualSonics, Canada), with animals maintained on a heated Physio Plate for continuous monitoring of body temperature, ECG, and respiration. Parasternal long- and short-axis views were acquired in B- and M-mode. Left ventricular internal diameter, ejection fraction, and anterior and posterior wall thickness were measured from the midventricular short-axis view using VevoLab software (version 5.5.0), with measurements obtained from at least 3 cardiac cycles and averaged.

#### 1.2. *In vivo* NRF2 activation

The *in vivo* NRF2 activation study was performed as previously described (ElBeck, Hossain et al. 2024). Briefly, *Mlp*$^{-/-}$ male and female mice were treated with the NRF2 activator AZ925 or a scramble control, and cardiac function and myocardial molecular responses were assessed. Echocardiographic measurements and RNA-sequencing data for *Hmox1*, *Nqo1*, and *Osgin1* were obtained from the previously published study. For the present study, myocardial tissue from the same experimental cohort was additionally analyzed by quantitative PCR to determine miR-208b-3p expression.

### 2. Human myocardial and public transcriptomic datasets

Multiple newly generated and publicly available transcriptomic datasets were used for comparative analysis of human cardiomyopathy and *Mlp*$^{-/-}$ myocardium. RNA-sequencing of myocardium from *in vivo* miR-208b-3p antagomir experiment and of miR-208b-3p-overexpressing HEK293 cells, as well as small RNA-sequencing of *Mlp*$^{-/-}$ and wild-type myocardium, was performed as part of the present study. Human myocardial RNA-sequencing data were obtained from Sielemann, Elbeck et al. (2020), Sweet et al. (2018; GEO accession GSE116250), and van Heesch et al. (2019). Human small RNA-sequencing data were obtained from Akat et al. (2014; GEO accession GSE53080). Previously published RNA-sequencing data from *Mlp*$^{-/-}$ myocardium were obtained from ElBeck et al. (2024; SRA accession PRJNA821374). Data analysis is described in the Data Analysis section below.

In the processed RPKM expression data that were obtained from the Gene Expression Omnibus (GEO; accession GSE116250), some *MYH7* values were coded as 999999999 in the processed dataset. These values were treated as invalid and excluded. Samples containing these values were retained in the overall study cohort but omitted from analyses requiring a valid *MYH7* expression measurement.

### 3. *In vitro* cell culture and experimental treatments

#### 3.1. Neonatal rat cardiomyocyte isolation, culture and treatments

Primary neonatal rat cardiomyocytes (NRCMs) were isolated from hearts of 3-day-old Sprague-Dawley rats. Following decapitation, hearts were rapidly collected and placed in ice-cold preservative buffer. The ventricles were separated from the atria, minced into small pieces, and enzymatically dissociated at 37°C using a gentleMACS Octo Dissociator with Heaters and the Neonatal Heart Dissociation Kit (Miltenyi Biotec), according to the manufacturer's instructions. The resulting cell suspension was diluted with cardiomyocyte plating medium consisting of high-glucose DMEM (4:1) supplemented with 5% horse serum, 2.5% fetal bovine serum, 20 mmol/L HEPES, and 100 U/mL penicillin/streptomycin. Cell suspensions were passed through 70-µm MACS Smart Strainers, followed by red blood cell lysis. To enrich for cardiomyocytes, the cells were pre-plated for 75 minutes at 37°C to allow preferential attachment of non-cardiomyocytes. The non-adherent cardiomyocyte-enriched fraction was collected, counted, and plated on 0.15% gelatin-coated 24-well plates at 3.33 × 10^5 cells per well. Cells were maintained at 37°C in a humidified incubator containing 5% $CO_2$. After 24 hours, and before redox-modulating experiments, cultures were rinsed once with PBS to remove nonviable cells and maintained for a further 48 hours in cardiomyocyte maintenance medium containing 2.5 µM of the mitotic inhibitor cytarabine.

Oxidative and redox-modulating experiments in primary neonatal rat cardiomyocytes (NRCMs) were performed as previously described (ElBeck, Hossain et al. 2024). Briefly, NRCMs were treated with hydrogen peroxide ($H_2O_2$), sulforaphane (SF), or N-acetyl-L-cysteine (NAC) to modulate intracellular redox status. $H_2O_2$ was prepared at 25, 50, 100, 200, or 300 µM and applied for 24 h, after which the medium was replaced with $H_2O_2$-free maintenance medium for an additional 24 h. Cells were collected at 6, 24, or 48 h from the start of treatment, as indicated. For SF and NAC experiments, NRCMs were treated with 5 µM SF, 3 mM NAC or combined for 48 h, with fresh treatment medium containing SF, NAC or both supplied every 24 h. For RNA expression analyses, cells were collected at 6, 24, or 48 h after treatment initiation by lysing them with TRIzol for RNA extraction. These concentrations SF and NAC were selected based on previously described dose-response experiments in NRCMs (ElBeck, Hossain et al. 2024). Cytarabine (2.5 µM) was included in the cardiomyocyte maintenance medium throughout all experimental conditions, except for the initial 24h culturing after isolation.

#### 3.2. *In vitro* miR-208b-3p gain-of-function studies

MiR-208b-3p gain-of-function experiments were performed using complementary approaches to increase intracellular miR-208b-3p levels. Cells were transfected with either a synthetic miR-208b-3p mimic or a miR-208b-3p expression construct, as specified for each experiment.

**Transfection with synthetic miR-208b-3p mimic:** HEK293 cells were grown in complete culture medium at 37°C in a 5% $CO_2$ incubator. Cells were seeded in 48-well plates and allowed to reach 70–80% confluency prior to transfection. Before transfection, cells were washed with PBS and incubated in 100 µl of Opti-MEM™ reduced serum (#31985062, Gibco) for 1 h to starve the cells. Transient transfections were performed using Lipofectamine RNAiMAX Reagent (Thermo Fisher Scientific) according to the manufacturer's instructions. Cells were transfected with either hsa-miR-208b-3p miRCURY LNA miRNA Mimic (QIAGEN, Cat. No. 339173, Product ID YM00472072-ADA) or the non-targeting miRCURY LNA miRNA Mimic Negative Control (QIAGEN, Cat. No. 339173, Product ID YM00479902-ADA) at a final concentration of 5, 100 and 200 nM. Lipid-RNA complexes were prepared in Opti-MEM Reduced Serum Medium and applied directly to the cells. Following incubation for 48 hours, the cells were harvested by lysing them with 75 + 75 µl of TRIzol (Invitrogen) for extracting the RNA.

***In vitro* transient overexpression of miR-208b-3p:** HEK 293 cells were plated in 12-well-plates (Corning) with the growth medium (DMEM, 10% FBS, Pen/Strep). Plated cells at a confluency of 70% were transfected with a mammalian shRNA knockdown vector carrying a hsa-miR-208b-3p. The expression of the small RNA was driven by the U6 promoter. A transcript of enhanced green fluorescent protein (EGFP) was added to the construct under the human phosphoglycerate kinase 1 promoter to visualize transfected cells by green florescence. For scramble control, cells were transfected with a mammalian gene expression vector carrying an enhanced green fluorescent protein (EGFP) transcript under the human eukaryotic translation elongation factor 1 α1 promoter (Table 2). This was the same control vector that was utilized in study-III.

**Table 2: Sequences of ShRNA utilized in the current work**

| Construct ID | Target gene | ShRNA sequences | Map |
|---|---|---|---|
| **VB201215-1116kpk** | hsa-miR-208b-3p | ACAAACCTTTTGTTCGTCTTAT | https://en.vectorbuilder.com/vector/VB201215-1116kpk.html |
| **VB180823-1120qay** | Scramble, EGFP | - | https://en.vectorbuilder.com/vector/VB180823-1120qay.html |

The transfection was performed with Lipofectamine™ 3000 reagents (Invitrogen) according to the manufacturer's instructions. In short, cells were washed with PBS and incubated in 400 µl of Opti-MEM™ reduced serum (#31985062, Gibco) for 1 h to starve the cells. Then an additional 100 µl of Opti-

MEM™ reduced serum was added, containing the lipofectamine transfecting reagents and 250 ng of the plasmid currying the miR-208b-3p or scramble constructs. Next day, 500 µl of fresh growth medium was added to each well. Thereafter, the growth medium was replaced daily with a fresh medium until day 5, when EGFP expression was examined under a florescence microscope, and then the cells were washed twice with PBS, and lysed with 90 + 60 µl of TRIzol (Invitrogen) for extracting the RNA.

Negative controls consisting of untransfected cells were simultaneously treated with the same procedure as those cells that were transfected with the plasmid carrying miR208b-3p construct, but without having any DNA in the transfection reagents.

### 4. Gene expression analysis

#### 4.1. Quantitative PCR

RNA was extracted with TRIzol (Invitrogen) following the manufacturer instructions, but adapting to smaller volumes. Extracted RNA was quantified with Qubit (Thermo Fisher Scientific). For the first strand synthesis of cDNA, SuperScript IV First-Strand Synthesis System (Invitrogen) was utilized with poly-T primers. The qPCR analyses were performed utilizing Universal SYBR Green Supermix (BioRad). All primers utilized in the assays are shown in Table 3 below.

**Table 3: qPCR primers utilized in the study**

| Gene | Forward primer | Reverse primer | Species |
|---|---|---|---|
| ***IDH2*** | GCGAAGCCCGTGGTGGAGAT | AGTCTGGTCACGGTTTGGGA | Human |
| ***MED13*** | GTGAAGAGCATATCACCCTTGC | AATGCCTGTCCTGTGAGAGTG | Human |
| ***PURB*** | CGATGTGGGCTGCAACAAATA | GTGATGGCATTGCGGTAGGA | Human |
| ***CBX1*** | GCCGGAGCGGATTATTGGAG | GTGGGCACTTGACATTGGC | Human |
| ***SP3*** | GTGATACCACAGATCCAGTCAGC | AGGTTTGATTAGAGCCAGGAATG | Human |
| ***SOX6*** | TACCTCTACCTCACCACATAAGC | ACATCGGCAAGACTCCCTTTG | Human |
| ***Myh6*** | CCAACACCAACCTGTCCAAGT | AGAGGTTATTCCTCGTCGTGCAT | Mouse |
| ***Myh7*** | CTCAAGCTGCTCAGCAATCTATTT | GGAGCGCAAGTTTGTCATAAGT | Mouse |
| ***Hmox1*** | CACGCATATACCCGCTACCT | CCAGAGTGTTCATTCGAGCA | Mouse |
| ***Osgin1*** | CCTCCGGTATCTGCCTGTC | GGAAAGGTACTCTAGGTCCTGG | Mouse |
| ***Idh2*** | GGAGAAGCCGGTAGTGGAGAT | GGTCTGGTCACGGTTTGGAA | Mouse |
| ***Hmox1*** | GAAGAAGATTGCGCAGAAG | GAAGGCGGTCTTAGCCTCTT | Rat |
| ***Nqo1*** | CTCGCCTCATGCGTTTTTG | CCCCTAATCTGACCTCGTTCAT | Rat |
| ***Osgin1*** | CCTCGGGTATCTGTCTATC | GGAAAGGTACTCTAGGTCCTGA | Rat |

For the synthesis of the cDNA of miR-208b-3p and U6 snRNA, TaqMan™ Fast Advanced Master Mix (#4444557, ThermoFisher Scientific) was utilizes according to the manufacturer's instructions, with the flowing thermal program (reverse transcription at 16°C for 30 min and then 42°C for 30 min, stop reaction 85°C for 5 min and then hold at 4°C).

For quantitative analysis of miR-208b-3p, the TaqMan™ MicroRNA assays; hsa-miR-208b (#4427975, Assay ID 002290) and U6 snRNA (#4427975, Assay ID, 001973) from Thermo Fisher Scientific were utilized, according to the manufacturer's instructions. The thermal program was (UNG activation at 50°C for 2 min, then enzyme activation at 95°C for 10 min, 40 cycles of [denature at 95°C for 15 sec, then Anneal /Extend 60°C for 60 sec, plate read].

Both cDNA synthesis and subsequent qPCR analyses of miRNA were performed in 384-well-plates utilizing CFX384 Touch™ Real-Time PCR Detection System (BioRad, US).

Echocardiographic data used to assess the correlation between left ventricular ejection fraction and myocardial miR-208b-3p expression in untreated male and female *Mlp*$^{-/-}$ mice and their wild-type (WT) littermate controls were obtained from ElBeck et al. *(ElBeck, Hossain et al. 2024)*. miR-208b-3p expression levels were measured as part of the present study.

#### 4.2. - Small RNA sequencing

Enriched small RNA fractions were extracted from the myocardia of 3 *Mlp*$^{-/-}$ mice and 3 WT littermate controls using mirVana™ miRNA Isolation Kit, following the manufacturer's instructions. The quantity of enriched small RNA fraction was estimated by Qubit™ through utilizing the RNA High Sensitivity Kit (Thermo Fisher Scientific), and the quality of the RNA was assessed through utilizing the Bioanalyzer Agilent Small RNA Kit. Small RNA libraries were prepared by utilizing TruSeq® Small RNA Library Prep Kit, and subsequently sequenced on HiSeq (rapid) Flowcell (1x50 bp reads) to yield ~15M reads per sample. Raw sequencing data were bioinformatically processed using a standard method to obtain a list of differentially expressed miRNAs.

#### 4.3. Sequencing of mRNA

Total RNA was extracted from the myocardium of *Mlp*$^{-/-}$ mice treated with miR-208b-3p antagomir or scramble control using TRIzol (Invitrogen), as described above. mRNA libraries were prepared using the Smart-seq2 protocol (Picelli, Faridani et al. 2014), with eight technical replicates per sample. HEK293 cells transfected with the miR-208b-3p expression vector (see In Vitro miR-208b-3p Gain-of-Function Studies) were processed using the same protocol, with a single sequencing replicate per sample.

### 5. Transcriptomic analysis

#### 5.1. Reads processing of RNA sequencing

RNA-sequencing reads were quality-controlled and processed using the bcbio pipeline (version 1.1.7-b). Reads were aligned using STAR (version 2.6.1d) to the mouse reference genome (mm10) or human reference genome (GRCh38), as appropriate, using Ensembl genome annotation release 91. Gene-level read counts were generated using featureCounts (version 1.6.4). For the *in vivo* myocardial samples, data from eight technical replicates were combined before differential expression analysis. For HEK293 samples transfected with the miR-208b-3p expression vector, the differential gene expression was analyzed using DESeq2 (R package version 1.26.0).

#### 5.2. Small RNA-seq data processing and analysis

Human small RNA-seq count data were obtained from the Gene Expression Omnibus (GEO; accession GSE53080). Analysis was performed in R using the edgeR package. Adult, pre-transplant left ventricular samples were included, comprising 10 non-failing controls, 22 DCM, and 14 ICM samples. Lowly expressed miRNAs were filtered using filterByExpr(), followed by library-size normalization using the trimmed mean of M-values (TMM) method. Differential expression was assessed using an edgeR generalized linear model for DCM versus control and ICM versus control. Differentially expressed miRNAs were defined as FDR <0.05 and absolute log2 fold change ≥1.

MiRNA identifiers were harmonized across mouse and human datasets according to mature miRNA identity and arm to enable cross-species comparison. Historical miRNA nomenclature was converted to the corresponding 3p or 5p mature miRNA designation, and annotation suffixes were standardized. Differentially expressed miRNA sets from *Mlp*$^{-/-}$, DCM, and ICM myocardium were compared using set intersection and set-difference analyses in R. Log2 fold-change values for the resulting miRNA sets were assembled for visualization as a heatmap in GraphPad Prism.

ChatGPT (OpenAI) was used to assist with aspects of the computational workflow and the preparation of the analytical methods description for this small RNA-seq analysis. The authors reviewed and verified the code and analytical outputs and take full responsibility for the accuracy and interpretation of the results.

### 6. Pathway, upstream regulator, and network analysis

Differentially expressed gene sets were analyzed using Ingenuity Pathway Analysis (IPA; QIAGEN) for pathway enrichment and upstream regulator analysis. For Figure 1, differentially expressed gene lists from human dilated cardiomyopathy (DCM) and *Mlp*$^{-/-}$ mouse myocardium were compared to identify shared and dataset-specific differentially expressed genes, which were visualized using a Venn diagram. Genes were selected using the criteria $p < 0.05$ and adjusted $p < 0.01$. The intersecting gene lists used for IPA analyses were generated in Microsoft Excel using the ¨vlookup¨ function to identify genes shared between the specified gene sets.

IPA pathway enrichment was used to identify biological pathways overrepresented among the selected genes. Enrichment significance was evaluated using the right-tailed Fisher's exact test. Where applicable, gene-selection thresholds were adjusted to accommodate IPA input limits, applying consistent criteria across datasets being compared. Predicted pathway activation or inhibition was assessed using IPA activation $z$-scores.

Upstream regulator analysis was performed to identify candidate regulators whose known downstream targets showed expression changes consistent with predicted regulator activation or inhibition. Predictions were based on the direction of observed gene-expression changes and the documented regulatory relationships between upstream regulators and their target genes. Predicted activation states were evaluated using IPA activation *z*-scores and were interpreted as computational predictions rather than direct measurements of regulator activity.

Network analysis present in Figure 1B was constructed by selecting pathways focusing on stress-response pathways associated with endoplasmic reticulum (ER) stress, protein homeostasis, and autophagy from the IPA enrichment results shown in Figure 1A. Selected pathways and associated molecular entities were connected using the Connect function in Ingenuity Pathway Analysis (IPA; QIAGEN) to generate a network illustrating relationships among these stress-response processes. Transcriptomic data from $Mlp^{-/-}$ myocardium were overlaid onto the network to visualize the expression patterns of the genes represented and the predicted activation state of these pathways. For Figure 5E, upstream regulator analysis was performed in IPA using transcriptomic data from $Mlp^{-/-}$ mice treated with miR-208b-3p antagomir to predict regulators associated with the observed gene-expression changes. Top stress-responsive regulators were selected, and their relationships were connected using the Connect function in IPA to generate a regulatory network. For Figure 6A, differentially expressed genes from miR-208b-3p-overexpressing HEK293 cells were mapped onto curated oxidative stress and NRF2-associated signaling networks in IPA. Previously reported miR-208b-3p target genes were also overlaid to examine their reported functional relationships with components of oxidative stress and apoptosis-associated signaling. The Connect function was used to visualize relationships among the selected molecular entities.

**7. Redox and Oxidative Stress Measurements**

Myocardial reduced glutathione (GSH), oxidized glutathione (GSSG), and malondialdehyde (MDA) were quantified by targeted liquid chromatography–tandem mass spectrometry (LC–MS/MS). Sample extraction and derivatization were performed as described previously (ElBeck et al., 2024), with modifications to the subsequent mass-spectrometric analysis.

For GSH and GSSG measurements, approximately 5 mg of powdered frozen myocardial tissue was mixed with 100 µL of aqueous extraction solution containing 5 mM N-ethylmaleimide (NEM), 0.05 mM deuterated GSH (GSH-d5) (Toronto Research Chemical, G597953), and 4 nM isotopically labelled GSSG internal standard ($^{[13]}C_4{}^{[15]}N_2$) (Toronto Research Chemical,

G597972). The two isotopically labelled internal standards were used to account for potential artifactual oxidation of GSH and reduction of GSSG during sample processing.

Samples were homogenized at 4°C using a TissueLyser, with three freeze–homogenization cycles, and incubated at room temperature for 20 min to facilitate GSH derivatization. Following centrifugation, an aliquot was retained for protein quantification, and proteins were precipitated by adding 400 μL methanol. After centrifugation, the supernatant was collected and dried under vacuum.

For MDA measurement, 1–2 mg of powdered frozen myocardial tissue was mixed with 50 μL of 1 M NaOH and heated at 60°C for 30 min. After cooling and sonication, an aliquot was retained for protein quantification. Deuterated MDA internal standard and trichloroacetic acid were added to precipitate proteins. The supernatant was reacted with 2,4-dinitrophenylhydrazine (DNPH) for 10 min at room temperature. The reaction was stopped by adding NaOH, and the resulting MDA-DNPH derivative was extracted twice using cyclohexane (1:1, v/v). The organic extracts were dried under vacuum.

After evaporation, samples were reconstituted with 0.1% of formic acid in water (200 μL for GSH/GSSG and 100 μL for MDA). Aliquots of the resulting supernatant (180 μL for GSH/GSSG; 100 μL for MDA) were transferred to amber LC-MS vials fitted with 300 μL glass inserts, capped, and loaded into the autosampler for UHPLC-MS analysis. ACQUITY Premier UPLC system coupled to a tandem mass spectrometer was used. Analytes were separated on an ACQUITY Premier UPLC HSS T3 column (2.1 × 100 mm, 1.8 μm) using 0.1% formic acid in water and 0.1% formic acid in methanol as mobile phases A and B, respectively. Chromatographic gradients were optimized separately for GSH/GSSG and MDA. The column temperature was maintained at 35°C, the flow rate was 350 μL/min, and the injection volume was 0.8 μL. Mass spectrometry was performed in positive-ion mode, with a capillary voltage of 3.0 kV, a desolvation temperature of 600°C, a cone gas flow of 150 L/h, and a desolvation gas flow of 1,000 L/h. Quantification was performed using selected-reaction-monitoring transitions and the corresponding internal standards. For GSH-NEM, both chromatographic peaks arising from regioisomeric derivatization products were integrated together. Analyte concentrations were normalized to total protein content. LC-MS/MS analyses were performed at the Small Molecule Mass Spectrometry Core Facility, that receives funding from the Infrastructure Board at Karolinska Institutet.

### 8. Statistical Analysis

Statistical analyses and graphical representations were performed using GraphPad Prism. Data are presented as mean ± standard error of the mean (SEM), with individual data points shown where indicated. Comparisons between two independent groups were performed using unpaired two-tailed *t*-tests, whereas paired two-tailed t-tests were used for longitudinal measurements. A paired one-tailed *t*-test was used for the specified longitudinal echocardiographic comparison. Correlations between continuous variables were assessed using Pearson's correlation coefficient. Statistical tests are specified in the corresponding figure legends. A *P* value <0.05 was considered statistically significant.

### 9. Data and Code Availability

All data and code associated with this study will be made publicly available upon publication in a peer-reviewed journal.

### 10. References:

ElBeck, Z., M. B. Hossain, H. Siga, N. Oskolkov, F. Karlsson, J. Lindgren, A. Walentinsson, D. Koppenhofer, R. Jarvis, R. Burli, T. Jamier, E. Franssen, M. Firth, A. Degasperi, C. Bendtsen, R. I. Menzies, K. Streckfuss-Bomeke, M. Kohlhaas, A. G. Nickel, L. H. Lund, C. Maack, A. Vegvari and C. Betsholtz (2024). "Epigenetic modulators link mitochondrial redox homeostasis to cardiac function in a sex-dependent manner." Nat Commun **15**(1): 2358.

Picelli, S., O. R. Faridani, A. K. Bjorklund, G. Winberg, S. Sagasser and R. Sandberg (2014). "Full-length RNA-seq from single cells using Smart-seq2." Nature Protocols **9**(1): 171-181.

Smith, A. J., R. E. Clutton, E. Lilley, K. E. A. Hansen and T. Brattelid (2018). "PREPARE: guidelines for planning animal research and testing." Lab Animal **52**(2): 135-141.
